# A novel RORγt^+^ antigen presenting cell type instructs microbiota-dependent regulatory T cell differentiation and tolerance during early life

**DOI:** 10.1101/2022.02.26.481148

**Authors:** Blossom Akagbosu, Zakieh Tayyebi, Gayathri Shibu, Yoselin A. Paucar Iza, Deeksha Deep, Yollanda Franco Parisotto, Logan Fisher, H. Amalia Pasolli, Valentin Thevin, Rasa Elmentaite, Maximilian Knott, Saskia Hemmers, Lorenz Jahn, Christin Friedrich, Jacob Verter, Zhong-Min Wang, Marcel van den Brink, Georg Gasteiger, Thomas G. P. Grünewald, Julien C. Marie, Christina Leslie, Alexander Y. Rudensky, Chrysothemis C. Brown

**Author notes:** Correspondence to (A.Y.R) or (C.C.B). These authors contributed equally (Blossom Akagbosu, Zakieh Tayyebi).

## Abstract

Establishing and maintaining tolerance to self- or innocuous foreign antigens is vital for preservation of organismal health. Within the thymus, medullary thymic epithelial cells (mTECs) expressing *A*uto*I*mmune *Re*gulator, Aire, play a critical role in self-tolerance through deletion of autoreactive T cells and promotion of thymic regulatory T (Treg) cell development. Within weeks of birth, a separate wave of Treg cell differentiation occurs in the periphery, upon exposure to dietary and commensal microbiota derived antigens, yet the cell types responsible for the generation of peripheral Treg (pTreg) cells are not known. Here we identified a new class of RORγt^+^ antigen-presenting cells (APC), dubbed Thetis cells (TCs), with transcriptional features of both mTECs and dendritic cells (DCs), comprising 4 major sub-groups (TC I-IV). We uncovered a developmental wave of TCs within intestinal lymph nodes during a critical early life window, coincident with the wave of pTreg cell differentiation. While TC I and III expressed the signature mTEC nuclear factor Aire, TC IV lacked Aire expression and were enriched for molecules required for pTreg generation, including the TGF-β activating integrin αvβ8. Loss of either MHCII or Itgb8 expression by TCs led to a profound impairment in intestinal pTreg differentiation, with onset of intestinal inflammation. In contrast, MHCII expression by RORγt^+^ group 3 innate lymphoid cells (ILC3) and classical DCs was neither sufficient nor required for pTreg generation, further implicating TCs as the tolerogenic RORγt^+^ APC with an essential early life function. Our studies reveal parallel pathways for establishment of tolerance to self and foreign antigen within the thymus and periphery, marked by involvement of shared cellular and transcriptional programs.

## Main

Within the thymus, a distinct lineage of epithelial cells establishes tolerance to self-antigens through deletion of autoreactive T cells and promotion of thymic regulatory T (Treg) cell differentiation^1,2^. These functions of medullary thymic epithelial cells (mTECs) are mediated in part through the expression of Aire, which regulates the ectopic expression of tissue-restricted antigens (TRAs)^1^. Another major site of tolerance induction resides within intestinal lymphoid tissue where an infant’s developing immune system is exposed to a slew of new dietary components and colonizing microbes upon weaning. Establishing a harmonious host-microbiota relationship in this early life developmental window is critical to prevent later onset of immune-mediated disorders^3,4^. Central to the establishment of tolerance to intestinal microbes is the differentiation of naive T cells into peripherally generated Treg (pTreg) cells upon encounter with commensal-derived antigens^5–8^. Yet the antigen presenting cell (APC) that promotes pTreg cell differentiation is not known. The narrow time window for establishing intestinal immune homeostasis suggests the presence of a developmentally restricted tolerogenic APC within the neonatal intestinal niche.

### Antigen presentation by RORγt^+^ APCs is critical for pTreg induction in early life

Extra-thymic pTreg cells, distinguished from their thymic counterparts by expression of the orphan nuclear receptor RORγt, arise in the mesenteric lymph nodes (mLN) in response to commensal bacterial antigens and play a critical role in suppressing inflammatory immune responses against gut microbes^5,8^. In an intriguing symmetry, mice deficient in MHC class II-restricted antigen presentation by RORγt^+^ cells (*MHCII^ΔRORγt^*), develop severe intestinal inflammation due to a failure to establish tolerance to commensal bacteria^9^, suggesting a potential connection between RORγt^+^ APCs and RORγt^+^ pTreg cell generation. To address this possibility, we analyzed *MHCII^ΔRORγt^* mice at 3 weeks of age, when pTreg cells accumulate within the intestine^5,7^. We observed a dramatic reduction in RORγt^+^ pTreg cells within the mLN and colonic lamina propria, along with expansion of CD44^hi^ effector T (Teff) cells (**Fig. 1a-d, Extended data Fig. 1a**). At 8 weeks of age these mice exhibited a sustained, severe reduction in RORγt^+^ pTreg cells along with expansion of colonic T_H_17 cells (**Fig. 1e**, **Extended data Fig. 1b**), in line with previous studies demonstrating a prominent role for pathobiont-specific RORγt^+^ pTreg cells in suppressing inflammatory T_H_17 cells^8^. Histological analysis demonstrated severe colitis with marked inflammatory cell infiltrate, mucosal ulceration and crypt loss (**Fig. 1f,g**), confirming a critical role for RORγt^+^ APCs in preventing dysregulated intestinal immune responses.

**Figure 1.**
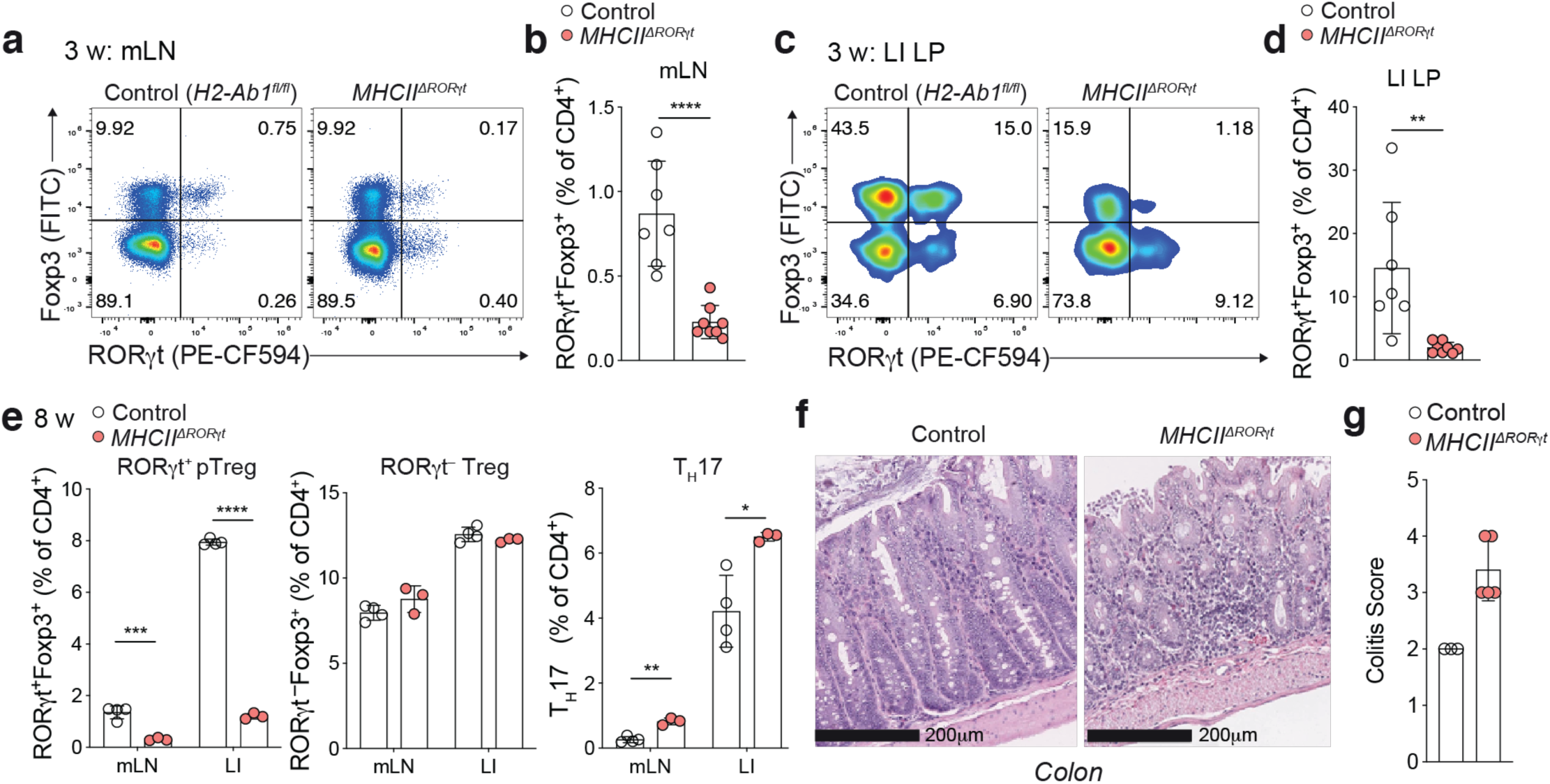
RORγt^+^ APCs promote pTreg differentiation and intestinal tolerance during early life. **a-d**, Representative flow cytometry of RORγt and Foxp3 expressing CD4^+^ T cell subsets and summary graphs for frequencies of pT_reg_ (RORγt^+^Foxp3^+^) cells in mesenteric lymph nodes (mLN) (a,b) and large intestine lamina propria (LI) (c,d) of 3 w old *MHCII^ΔRORγt^* (*n* = 7) and control (*H2-Ab1^fl/fl^*) (*n* = 8) mice from two independent experiments**. e**, Adult (8 week) *MHCII^ΔRORγt^* (*n* = 4) and control (*n* = 3) mice were analyzed for frequencies of pTreg (RORγt^+^Foxp3^+^), RORγt^−^ Treg cells, and T_H_17 (Foxp3^−^RORγt^+^) among CD4^+^ T cells in indicated tissues. **f** Histological analysis of H&E stained sections of the colon. Scale bars represent 200 μm. **g,** Histological colitis score in 12-week-old *MHCII^ΔRORγt^* (n=5) and control (n=3) mice. Error bars: means ± s.e.m. Each symbol represents an individual mouse. \**P* < 0.05; \*\**P* < 0.01; \*\*\**P* < 0.001; \*\*\**P* < 0.0001; unpaired two-sided *t*-test.

Given previous reports of a developmental window for intestinal immune tolerance^10,11^, we wondered whether the pTreg cell deficit in adult *MHCII^ΔRORγt^* mice reflected a failure to generate pTreg cells in early life or a continuous requirement for RORγt^+^ APC-instructed pTreg cell differentiation. We therefore generated a novel *Rorc^Venus-creERT2^* mouse for identification and temporal manipulation of RORγt^+^ cells (**Extended data Fig. 2a,b**). Analysis of *Rorgt^cre^R26^lsl-tdTomato^Rorc^Venus-creERT^*^2^ mice confirmed that expression of Venus protein, translated downstream of exon 11, faithfully reflected expression of the RORγt isoform within the mLN and large intestine (**Extended data Fig. 2c**). Surprisingly, continuous ablation of MHCII on RORγt^+^ APCs in adult *Rorc^Venus-creERT2^H2-Ab1^fl/fl^* mice treated with tamoxifen from 8-13 weeks of age, resulted in only a very modest decrease in pTreg cells within the mLN and LI (**Extended data Fig. 2d,e**), indicating minimal contribution of *de novo* pTreg cell differentiation to the pTreg cell pool of adult mice with stable microbial communities. Together these results demonstrate an essential role for an early life RORγt^+^ APC in pTreg cell generation and raise the question as to the nature of the tolerogenic RORγt^+^ APC.

### Identification of a novel lineage of RORγt^+^ antigen presenting cells

A number of candidate APC types have been suggested to regulate tolerance to the intestinal microbiota including dendritic cells (DCs) and MHCII^+^ ILC3 cells, also known as lymphoid tissue inducer (LTi)-like cells. Amongst these, loss of tolerance to commensals in *MHCII^ΔRORγt^* mice has previously been attributed to ILC3s on the assumption that they represent the only RORγt^+^MHCII^+^ cell type^9,12^. Since ILC3s do not prime T cells *ex vivo*, it was suggested that ILC3s mediate negative selection of commensal reactive Teff cells^9^. However, recent studies have identified RORγt expressing DCs^13,14^ as well as RORγt^+^Aire^+^ cells^15^, initially described as ‘ILC3-like’ cells but subsequently shown to more closely resemble DCs^16^. The role of these cells in immune tolerance remains unknown. Critically, the spectrum of RORγt^+^ APCs has not been examined within the mLN at the time when pTreg cells first arise. RORγt^+^ pTreg cells first appeared within the mLN between postnatal days 10 and 14 (P10–14) with rapid accumulation thereafter (**Extended data Fig. 3a**). We therefore performed paired single cell (sc)RNA/ATAC-sequencing of CD45^+^Lin^−^RORγt(Venus)^+^MHCII^+^ cells isolated from the mLN at 2 weeks of age (**Fig. 2a**, **Extended data Fig 3b**). After quality filtering, we retained chromatin accessibility and transcriptional profiles on 10,145 cells. Unsupervised clustering of either the RNA or ATAC-seq data revealed two major cell types (**Fig. 2b-d; Extended data Fig. 3c**). The first represented ILC3s spanning their full developmental spectrum including a RAG1^+^ ILC3 progenitor (ILC3p), proliferating and mature NCR^+^ ILC3s, and CCR6^+^ LTi cells (**Fig. 2d, Extended data Fig. 3c,d**). The second cell type did not express canonical ILC genes. This population, which was distinguished by an intriguing combination of both epithelial and DC-associated transcription factors and cell surface molecules, consisted of 4 subsets and a small cluster of proliferating cells (**Fig. 2b-d**, **Extended data Fig. 3c-e**). Although these cells expressed the DC marker *Zbtb46*, this transcript was also unexpectedly highly expressed by MHCII^+^ ILC3s, a finding confirmed by analysis of *Zbtb46^GFP^Rorgt^cre^R26^lsl-tdTomato^* mice (**Extended data Fig. 3f**). To elucidate the identity of non-ILC3 RORγt^+^ APCs, we compared similarity of pseudo-bulk transcriptomes across a comprehensive database of immune and stromal cells (ImmGen). As expected, ILC3 scRNA clusters aligned with ILC3s, whereas the remaining clusters exhibited surprisingly high correlation with both mTECs and DCs (**Fig 2e**), including specific expression of Aire, the signature mTEC transcription factor, within clusters I and III (**Fig 2f**). Independent cross-referencing of these cells with immune and epithelial cell atlases using CellTypist^17^, a machine learning tool for precise cell type annotation, also showed their overlapping transcriptional features with both DCs and a generic epithelial cell (**Extended data Fig. 3g,h**), consistent with their expression of p63, a critical regulator of epithelial cell differentiation^18,19^. In light of the ‘shape-shifting’ hybrid phenotype of this group of RORγt^+^ APCs, we refer to these cells as Thetis cells (TCs). Comparison of TC cluster identity defined by chromatin accessibility or gene expression revealed near perfect congruence (**Extended data Fig. 2f**), confirming that TCs comprise four distinct cell types (TC I-IV). Analysis of pseudo-bulk transcriptomes for TC sub-groups alongside published single-cell thymic epithelial transcriptomes^20^, demonstrated overlap with distinct clusters of mature mTEC subsets, in particular mature Aire^+^ (mTEC II) and ‘post-Aire’ (mTEC III) subsets (**Fig. 2g**). Overall, these data demonstrated the existence of a novel RORγt^+^ cell type, distinct from ILC3s, present within intestinal lymph nodes during early life.

**Figure 2.**
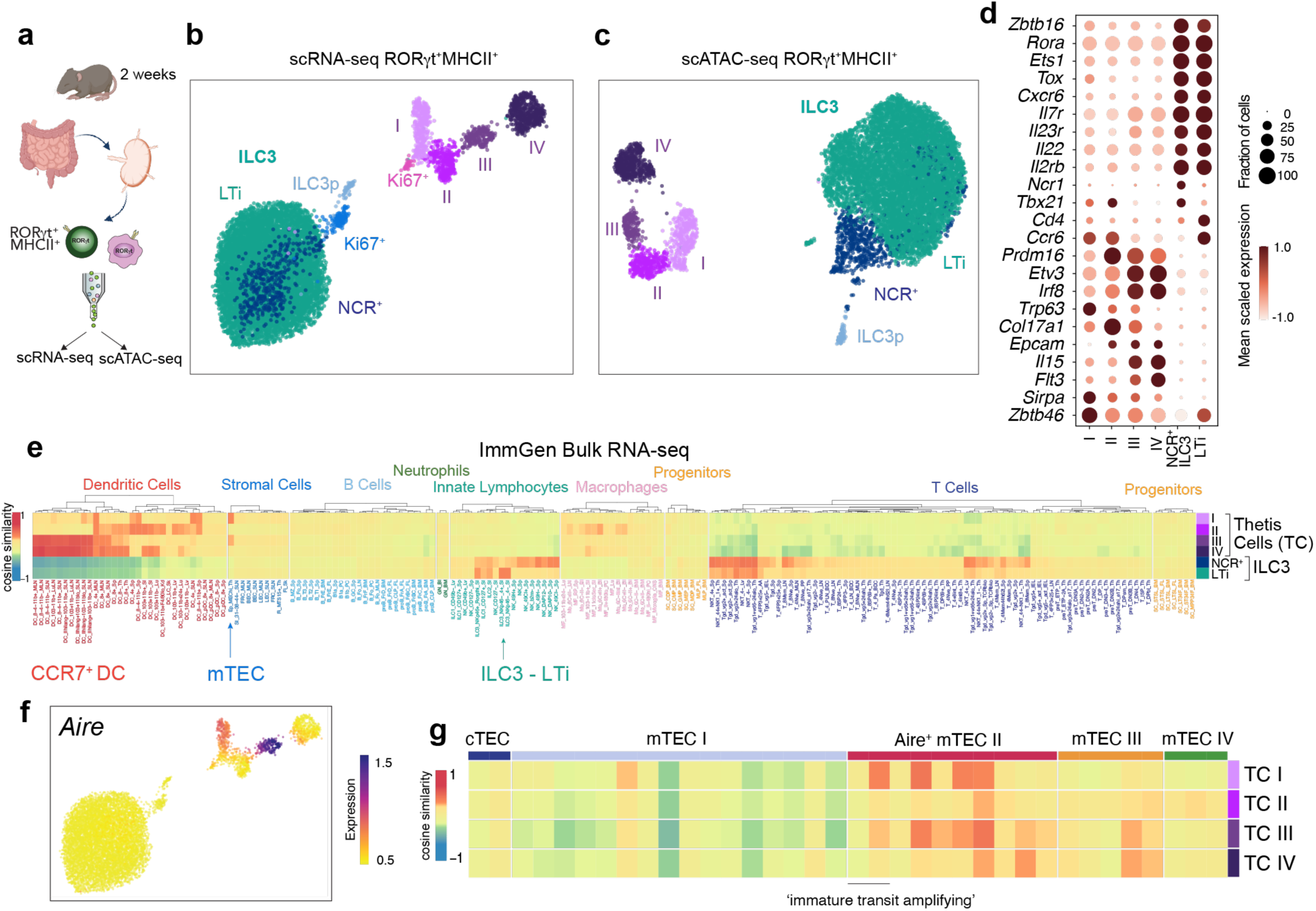
Identification of a novel RORγt^+^ antigen presenting cell lineage. **a**, Schematic of paired single cell transcriptome and epigenome profiling of Lin^−^RORγt^+^MHCII^+^ cells from mLN of 2-week-old *Rorc^Venus-^*^c*reERT2*^ mice (pooled from 16 biological replicates). UMAP visualization of 10,145 cells profiled by scRNA-seq (**b**) or scATAC-seq (**c**), colored by cluster annotation. **d**, Dot plot showing the expression of canonical ILC3 or cluster I-IV marker genes **e**, Similarity between cell types identified in (b) and ImmGen bulk microarray profiles for immune and stromal cells **f**, scRNA-seq UMAP colored by imputed expression of *Aire*. **g**, Similarity between TC subsets in (b) and mTEC subsets.

### Characterization of the phenotypic landscape of Thetis cells

Extra-thymic Aire expression has previously been reported in migratory CCR7^+^ DCs^21^. Of note, the gene expression signature that distinguishes CCR7^+^ DCs from their CCR7^−^ counterparts is not DC lineage defining, but rather represents a particular transcriptional program that can be acquired by classical DC subsets, cDC1 and cDC2, as well as other APC types^22,23^, reflecting enhanced cell migration, T cell priming capacity and expression of immune regulatory molecules^23^. The shared expression of Aire in TCs and CCR7^+^ DCs prompted us to examine the relationship between these two cell types. Analysis of *Rorc^Venus-creERT2^Aire*^GFP^ mice confirmed wide-spread Aire(GFP) expression by Lin^−^CXCR6^−^ CD11c^+^MHCII^+^CCR7^+^ cells encompassing both CCR7^+^ DC1 and DC2 (**Extended data Fig. 4a,b**); however, < 4% of CCR7^+^ cells expressed RORγt. Moreover, CXCR6^−^RORγt^+^MHCII^+^ TCs were also found amongst CCR7^−^ and CD11c^−^ MHCII^+^ cell populations (**Extended data Fig. 4a,b**), suggesting that TCs were distinct from CCR7^+^ DCs. To gain further insight into the distinguishing features of Aire-expressing TCs, DCs, and mTECs we performed orthogonal SMART-seq2 (SS2) scRNA-seq analysis of Lin^−^ RORγt^+^MHCII^+^ cells isolated from the mLN of 3-week-old *Rorc^Venus-creERT2^* mice, in parallel with mLN Aire^+^ DCs and Aire^+^ mTECs from age-matched *Aire*^GFP^ mice (**Extended data Fig. 4c-e**). Clustering analysis, combined with mapping of SS2 transcriptomes to the droplet 10X dataset, confirmed the presence of LTi-like ILC3 and TC I-IV (**Fig. 3a; Extended data Fig. 4f,g**). Within the SS2 dataset, Aire^+^ mTECs clustered with TC I (**Extended data Fig. 4f,g**). Nevertheless, a direct comparison revealed unique expression of epithelial genes (*Foxn1, Krt17, Krt8*), the thymic marker gene *Tbata*, and *Fezf2* by mTECs, whereas TC I expressed genes associated with DCs (*Ccr7*, *Cd83, Dpp4*) (**Extended data Fig. 4h, Supplementary Table 2)**. In addition, TCs exclusively expressed *Ptprc* (CD45) and RORγt (**Extended data Fig. 4i**), a finding confirmed by analysis of mTECs from *Rorc^Venus-creERT2^* mice (**Extended data Fig. 4j**). Despite overlapping markers, TCs clustered separately from Aire^+^ DCs (**Extended data Fig. 4f,g**), distinguished by a number of immune regulatory genes (**Extended data Fig. 4k, Supplementary Table 3**), underscoring the distinct identity of these two cell types. Furthermore, Aire protein expression was readily detectable in 10-15% of the TC population, but not DCs (**Fig. 3b**), consistent with the lower level of *Aire* transcript observed within DCs.

**Figure 3.**
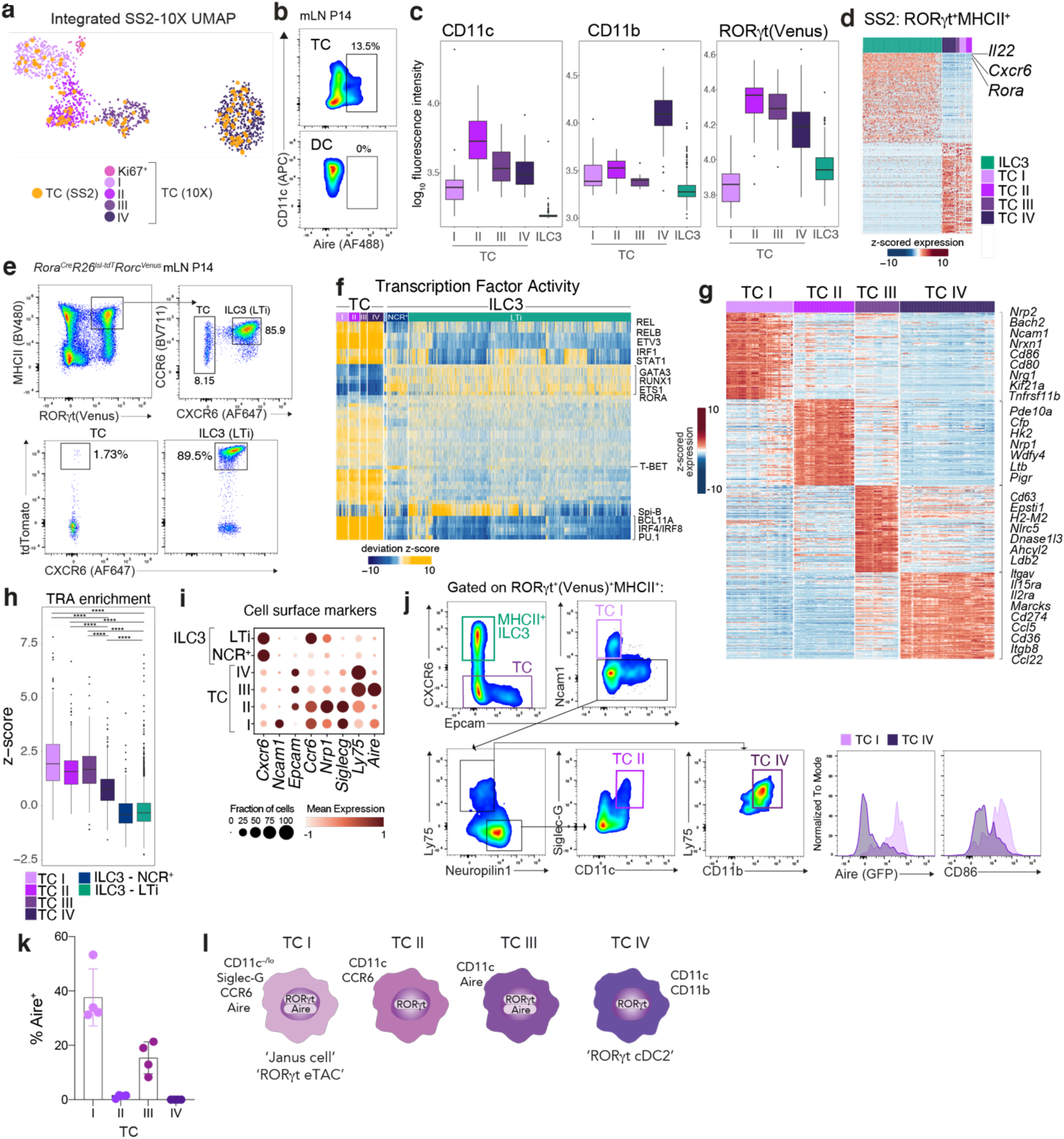
Transcriptional, epigenetic and ontological features of TC subsets. **a**, UMAP visualization of integrated 10X Genomics and Smart-Seq2 (SS2) scRNA-seq analysis for RORγt^+^MHCII^+^ TCs, colored by SS2 TC transcriptomes or 10X cluster annotation. **b**, Intracellular expression of Aire protein by TCs and DCs. **c**, Index-sorting summary graphs for CD11c, CD11b cell surface protein and Rorgt(Venus) fluorescence intensity. **d**, Heatmap showing expression of top differentially expressed genes between TCs and MHCII^+^ ILC3s profiled by SS2, identifying *Rora* as an ILC3/TC distinguishing gene. **e**, Representative flow cytometric analysis of tdTomato expression in MHCII^+^ILC3 and TCs isolated from mLN of *Rorc^Venus^Rora^cre^R26^lsl-tdTomato^* fate-mapped mice at P14*, n* = 4 mice. **f**, Heatmap reporting scaled chromVAR deviation TF motif scores (left) and corresponding TF gene expression values (right) for top TF gene-motif pairs in TCs in scATAC-seq data. **g**, Heatmap showing scaled, imputed expression of top 125 differentially expressed genes (one vs the rest, FC>1.5, adj. *P*<0.01) for each TC cluster. **h**, Enrichment of tissue-restricted antigen (TRA) genes across TC and ILC3 subsets identified in Fig. 1a. **i**, Dot plot showing expression of select cell surface markers, differentially expressed between TC subsets. **j,** Gating strategy for identification of TC subsets and expression of signature molecules. Plots are representative of 6 mice from two independent experiments. **k**, Intracellular expression of Aire protein by TC subsets; each symbol represents an individual mouse (*n* = 4). **l**, summary of TC I-IV phenotypes. Box plots (**c,h**) indicate the median (center lines) and interquartile range (hinges), and whiskers represent min and max, dots represent outliers. *P*<0.0001 ****; Mann Whitney U test.

Index sorting analysis of cell surface markers revealed that TCs spanned a spectrum from CD11c^−/lo^ (TC I) to CD11c^hi^ (TC II-IV) cells (**Fig. 3c, Extended data Fig 5a)**. In addition, TC IVs were distinguished by high levels of CD11b expression. These findings suggest that RORγt^+^CD11c^+^CD11b^+^ cells, previously identified amongst Tbet^−^ cDC2B^14^, represent TC IV. TC II-IV expressed higher levels of Venus(RORγt) than ILC3s (**Fig. 3c**), reflecting *Rorgt* promoter activity (**Extended data Fig. 5b**). Although TC I expressed lower levels of RORγt, all TC subsets expressed sufficient levels of *Rorgt* to drive *Rorgt*-cre-mediated recombination as evidenced by the near universal expression of tdTomato by CXCR6^−^ Venus(RORγt)^+^MHCII^+^ TCs isolated from *Rorc^Venus-creERT2^Rorgt^cre^R26^lsl-tdTomato^* mice (**Extended data Fig. 5c**). Of note, ILC3s, traditionally identified as CD90(Thy1)^+^ cells, encompassed both CD90^−^ and CD90^+^ cells and did not express CD11c (**Fig. 3c**, **Extended data Fig. 5d**). To assess whether the partially overlapping transcriptional features of TCs with DCs and mTECs were coupled to similar or distinct morphological attributes, we analyzed TCs by electron microscopy (EM). DCs, Aire^+^ mTECs, and MHCII^+^ ILC3 cells served as reference populations. Although CD11c^−^RORγt^+^Aire^+^ cells have previously been described as lymphoid^15^, EM analysis revealed that both CD11c^−^Aire^+^ TC I and CD11c^+^ TCs more closely resembled myeloid cells, in contrast to MHCII^+^ ILC3s which had a typical lymphoid appearance. TCs were distinct from classical CCR7^+^ and CCR7^−^ DCs, as well as mTECs, and featured distinctive mitochondria with rounded, condensed cristae **(Extended data Fig. 5e**). To further probe TC localization and morphology, we examined their spatial distribution in the mLN of Aire^GFP^ mice at P18. Immunofluorescence staining confirmed the presence of Aire^+^ TC (TC I and III) as well as Aire^−^CD11b^+^ TC IV with TC IV preferentially located within the paracortex (**Extended data Fig. 5f**). Analysis of mLN from *Rorc^Venus-creERT2^* mice confirmed similar spatial distribution of TC IV and RORγt^+^ pTreg cells (**Extended data Fig. 5g,h**).

### Thetis cell ontogeny

To address the ontogeny of TCs, we first analyzed DC fate-mapping *Clec9a^cre/wt^R26^lsl-tdTomato^Rorc^Venus-creERT2^*Aire^GFP^ reporter mice in which both cDC1 and cDC2 are labelled by virtue of Clec9a expression in DC progenitors^24^. In contrast to both Aire^+^ and Aire^−^ DC2s, < 5% of TCs were tdTomato^+^ (**Extended data Fig. 6a**), likely reflecting the small proportion of TCs with detectable *Clec9a* expression (**Extended data Fig. 3e**), rather than descendancy from a CLEC9A^+^ pre-DC. Given reports of a lymphoid pathway for cDC differentiation^25^, we next used a novel *Rag1^RFP-creERT2^R26^lsl^*^-YFP^ mouse model to fate-map progeny of lymphoid progenitor cells following neonatal 4-hydroxytamoxifen (4-OHT) administration. In contrast to labeling of T cells and ILC3s, YFP^+^ cells were absent amongst Lin^−^MHCII^hi^CXCR6^−^ cells encompassing TCs (**Extended data Fig. 6b**). These findings suggested that TCs are ontogenically and transcriptionally distinct from classical DCs. Thus, the overlapping phenotype of TCs and DCs, in particular CCR7^+^ DCs, likely reflects shared functions related to cell migration and antigen presentation rather than shared ontogeny (**Extended data Fig. 6c**). RORγt^+^Aire^+^ cells have previously been suggested to be related to ILC3s^15,26^; however, the absence of *Rag1* fate-mapped TCs suggested that TCs were not descended from the RAG1^+^ ILC3p identified within our scRNA-seq dataset (**Extended data Figure 3c**). Furthermore, TCs did not express RORα (**Fig. 3d**), a critical gene for ILC development, expressed by ILC precursors (ILCp) and mature ILC subsets^27–29^. The exclusive expression of RORα by MHCII^+^ ILC3s, allowed us to determine the lineage relationship between ILCp and TCs using *Rorc^Venus^Rora^cre^R26^lsl-tdomato^* fate mapping mice. Within the mLN, ∼90% of LTi cells and 70% of ILC1/NK cells were tagged with tdTomato vs <2% of TCs (**Fig. 3e, Extended data Fig. 6d**), confirming that TCs are not developmentally related to ILCs. To address the reverse relationship – TCs as the precursor to ILC3s – we analyzed single-cell transcriptional dynamics using single-cell RNA velocity^30^ to computationally define lineage relationships between TCs and ILC3s. This analysis identified established differentiation trajectories emanating from the ILC3p to NCR^+^ ILC3s and LTi cells (**Extended data Fig. 6e**). In contrast, no “connectivity” was observed between TCs and ILC3s, in either direction. Together these results demonstrate that TCs represent a new class of cells, distinct from both ILCs and classical DCs.

To determine the transcription factors (TF) that regulate TC differentiation and heterogeneity, we turned to our scATAC-seq data, integrating differential TF motif activity with gene expression. This analysis identified activity of canonical ILC3 TFs in ILC3s including RORα, GATA3 and TCF1, as well T-bet in NCR^+^ ILC3s (**Fig. 3f**), validating our approach. In contrast, TCs were distinguished by activity of a unique group of TFs including Spi-B, a critical regulator of mTEC differentiation, as well as core TFs governing myeloid cell differentiation (PU.1, BCL11A, IRF8 and IRF4) (**Fig. 3f**), in agreement with their transcriptional overlap with both mTECs and DCs. Notably, several of the signature TC TFs have been shown to regulate Aire expression in mTECs^31^, suggesting a conserved transcriptional network shared by these two cell types. Together these findings establish the unique identity of TCs, delineating their shared and distinct features with both mTECs and DCs

### TC subsets are enriched for distinct gene expression programs

To gain further insight into the functions of TC subsets, we examined their distinguishing transcriptional features (**Figure 3g**). TC I expressed canonical Aire^+^ mTEC markers, including *Aire*, *CD80*, *CD86*, *Tnfrsf11b* (OPG), and genes associated with neuronal adhesion, signaling and growth (*Nrxn1*, *Nrn1, Ncam1*). Of note, a recent study of peripheral Aire-expressing cells identified a population of ‘mTEC-like’ RORγt^+^Aire^+^ cells within lymph nodes, similarly distinguished by neuronal genes^16^, likely representing Aire^+^ TC I. TC II was distinguished by exclusive expression of several distinctive genes including *pIgR* and *Cldn7*, signature molecules for a group of mTECs with a history of Aire expression^32^, further highlighting parallels between mTEC and TC subsets. TC III expressed high levels of Aire as well as *Nlrc5*, a critical regulator of MHC Class I genes, suggesting unique roles in antigen presentation. TC IV expressed immune-regulatory genes (*Cd274*) as well as genes associated with cell migration (*Marcks*, *Cxcl16).* Given the critical role of Aire in promiscuous expression and presentation of tissue-restricted antigens (TRAs)^33^, we analyzed TRA expression levels amongst TCs in parallel with mTECs. This revealed significant enrichment of TRA genes in TCs, relative to MHCII^+^ ILC3s, most pronounced amongst TC I-III (**Fig. 3h**) as well as expected enrichment of TRAs in Aire^+^ mTECs relative to other thymic epithelial cell subsets (**Extended data Fig. 7a**). TRA abundance was similar across mTECs and TCs, with stochastic expression observed in both cell types (**Extended data Fig. 7b-d**), consistent with previous patterns of TRA expression reported in mTECs^20^. TC-enriched TRAs spanned antigens unique to organs that are specifically affected by autoimmune inflammation in Aire deficiency, including pancreas, testis, adrenal and liver (**Extended data Fig. 7e, Supplementary Table 3**), further highlighting convergence of mTECs and TCs in relation to their transcriptional features and subset composition. Analysis of differential chromatin accessibility and motif enrichment across TC subsets suggested several subset-specific transcriptional regulators, further underpinning the observed TC heterogeneity (**Extended data Fig. 7f**).

To validate the observed TC phenotypes, we devised a panel of markers for flow cytometry (**Fig. 3i**). MHCII^hi^RORγt^+^ ILC3s were distinguished from TCs by expression of CXCR6 (**Fig. 3i,j**). We confirmed the presence of TC subsets expressing signature cell surface markers with the expected pattern of Aire expression in TC I and III (**Fig. 3j,k**). Of note, CCR6 and Siglec-G, suggested to be markers for RORγt^+^Aire^+^ cells^15,34,35^, were expressed by TC I-II but not TC III-IV (**Fig. 3i, Extended data Fig. 7g**). In addition, TCs spanned IL7R^neg-hi^ cells (**Extended data Fig. 7h**). Of note, IL7Rα was also expressed by CCR7^+^ DCs (**Extended data Fig. 7i**). Moreover, a recent report demonstrated IL7R ‘fate-mapping’ in ∼25% of cDCs^36^, precluding this approach as a means of lymphoid cell lineage tracing. Thus, while Aire^+^ TC I represent RORγt^+^Aire^+^ CCR6^+^/Siglec-G^+^ cells (referred to either as ‘Janus cells’ or RORγt^+^ extrathymic Aire-expressing cells (eTACs) in two concurrent analyses of RORγt^+^ APCs^34,35^), TC II-IV described herein extend the spectrum of non-ILC3 RORγt^+^MHCII^+^ cells beyond Aire-expressing cells (**Fig. 3l**). Together, our analyses suggest that TC subsets are molecularly and functionally distinct and point to a role for TC IV in pTreg differentiation.

### Antigen presentation by ILC3s or DCs is not required for intestinal tolerance

Given the overlapping phenotype of TCs with professional APC types with known roles in T cell tolerance, we hypothesized that TCs were the relevant RORγt^+^MHCII^+^ cell type for instructing pTreg cell differentiation. A direct comparison of TC and ILC3 transcriptomes, as well as cell surface protein expression, confirmed that TCs were enriched for molecules associated with antigen presentation, T cell activation and cell migration, in contrast to MHCII^+^ ILC3 cells (**Fig. 4a,b, Extended data Fig. 8a**). Furthermore, in contrast to TCs, we did not observe CCR7 protein expression on MHCII^+^ ILC3s (**Fig. 4c**), despite detectable *Ccr7* transcript, making their migration from large intestine to mLN unlikely. To examine the antigen-presenting ability of TCs, we analyzed cell-surface I-A^b^ bound CLIP peptide on *H2-Dma*^−/−^ or littermate wild-type TCs, which confirmed efficient H-2M mediated CLIP displacement (**Extended data Fig. 8b**). Accordingly, staining with the 25-9-17s monoclonal antibody (mAb), which binds to specific non-CLIP peptide-I-A^b^ complexes^37^, demonstrated equivalent levels of expression by TCs and classical DCs (**Extended data Fig. 8c**). To further examine MHC class II antigen presentation by TC we bred *Rorc^Venus-creERT2^* with BALB/c mice and confirmed expression of an endogenously processed self-peptide, Eα bound I-A^b^, using the YAe mAb (**Extended data Fig. 8d**). To address the ability of TCs to induce pTreg cells *ex vivo*, we co-cultured TC subsets (either CCR6^+^ TC I and II or CCR6^−^ TC III and IV) with naïve C7 TCR transgenic CD4^+^ T cells and their cognate peptide under suboptimal Treg-inducing conditions. Strikingly, TCs demonstrated significantly greater ability to promote Treg differentiation relative to cDC2s with greatest efficacy observed amongst TC III and IV (92.7% TC III/IV vs 39.4% Foxp3^+^ cDC2, respectively; **Extended data Fig. 8e**). Together these results suggest that TCs are competent APCs.

**Figure 4.**
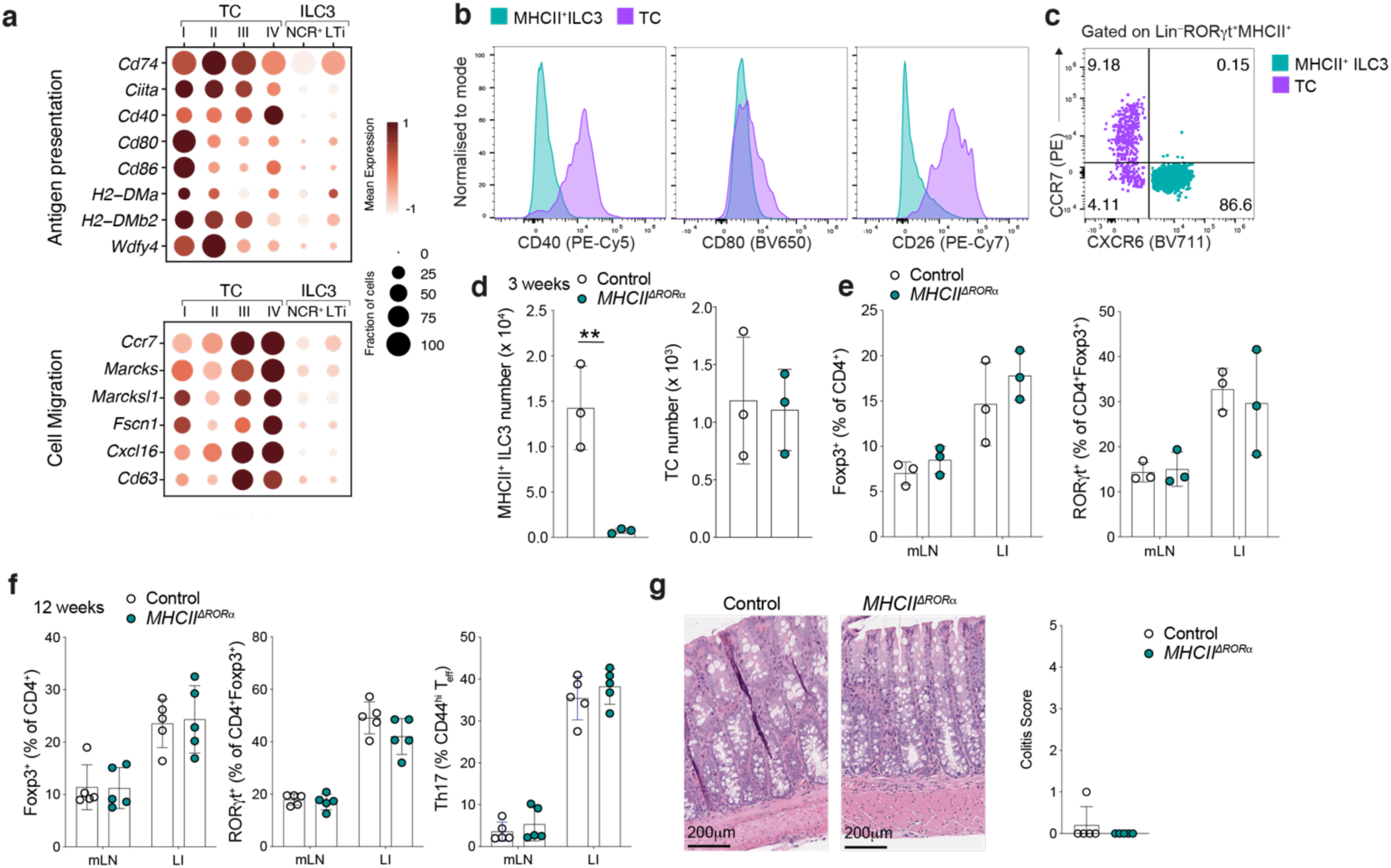
Antigen presentation by ILC3s is not required for intestinal pTreg differentiation. **a**, Dot plot showing expression of genes related to antigen presentation, T cell priming and cell migration across TC and MHCII^+^ ILC3 clusters (Fig. 2b). **b**, Flow cytometry of mLN ILC3s (CXCR6^+^RORγt^+^MHCII^+^) and TCs (CXCR6^−^RORγt^+^MHCII^+^) showing expression of indicated chemokine receptors, co-stimulatory and immune-regulatory molecules by TC and ILC3 in mLN of P18 *Rorc^Venus-creERT2^*Aire^GFP^ mice. **d**-**e**, Immune cell composition of 3-week-old *MHCII^ΔRORα^* (*n* = 3) and *Rora*^cre^*H2-Ab1^fl/wt^* (*n* = 3) mice. (**d**) Number of MHCII^+^ ILC3s and TCs in mLN. (**e**), Frequency of total T_reg_ (Foxp3^+^) and RORγt^+^ pTreg cells. **f**, Frequency of total T_reg_ (Foxp3^+^), RORγt^+^ pTreg cells and Th17 cells in mLN and large intestine lamina propria (LI) of 12w *MHCII^ΔRORα^* (*n* = 5) and *Rora*^cre^*H2-Ab1^fl/wt^* (*n* = 5) mice. **g,** Representative histological analysis of H&E stained sections of the colon for mice in (f) and summary histological colitis score. Scale bars represent 200 μm. Error bars: means ± s.e.m. Each symbol represents an individual mouse. *P* = non-significant; unpaired two-sided *t*-test.

Our previous analysis demonstrated the utility of *Rora*^cre^ to selectively target ILC3s but not TCs. Importantly, ILC3s were the only MHCII^+^ RORα fate-mapped cell type within the mLN (**Extended data Fig. 8f**), establishing *MHCII^ΔRORα^* mice as a genetic model for studying the functional role of ILC3 antigen presentation. Indeed, analysis of 3-wk-old *MHCII^ΔRORα^* mice confirmed a complete loss of MHCII expression on ILC3s, with no changes in TCs (**Fig. 4d**, **Extended data Fig. 8g**). Surprisingly, the intestinal T cell composition was not perturbed in *MHCII^ΔRORα^* mice, with equivalent proportions and numbers of CD4^+^ Teff and Treg cells, including RORγt^+^ pTreg cells, within the mLN and large intestine (**Fig. 4e, Extended data Fig. 8h,i**). To exclude a role for ILC3s in pTreg differentiation in later life, we examined *MHCII^ΔRORα^* mice at 12 weeks of age. In contrast to *MHCII^ΔRORgt^* mice, *MHCII^ΔRORα^* adult mice had normal pTreg frequencies with no evidence of altered T cell activation states (**Fig. 4f**) and lacked histological signs of colonic inflammation (**Fig. 4g**), further confirming that MHCII-mediated antigen presentation by ILC3s is not required for intestinal tolerance. Of note, our earlier analysis identified exclusive expression of IL-22 by ILC3s (**Fig. 3d**). However, consistent with the low frequency of fate-mapped MHCII^+^ILC3s in *Il22^cre^Rosa^lsl-tdTomato^* mice (<5%; **Extended data Fig. 8j**), *MHCII^ΔIL22^* mice exhibited only minimal loss of MHCII expression by ILC3s with no impact on the pTreg cell population (**Extended data Fig. 8k**). Besides ILC3s, antigen presentation by sub-immunogenic DCs is thought to favor T cell tolerance. Although TCs may have been inadvertently targeted by studies utilizing “DC-specific” Cre drivers^38^ due to expression of both CD11c and Zbtb46, the absence of Clec9a fate-mapped TCs allowed us to revisit a role for classical DC in pTreg differentiation through analysis of *Clec9a^cre/cre^H2-Ab1^fl/fl^* (*MHCII^ΔDC^*) mice in which DCs but not TCs were rendered MHCII-deficient (**Extended data Fig. 8l**). Surprisingly, we did not observe changes in RORγt^+^Foxp3^+^ cells in these mice, (**Extended data Fig. 8m**). Overall, these findings demonstrate that MHCII antigen presentation by ILC3s or cDCs is dispensable for pTreg cell differentiation, leaving TCs as the pTreg-inducing RORγt^+^ APC.

### A developmental wave of TCs during early life induces pTreg cells in an Itgb8 dependent manner

Given the narrow temporal window of opportunity for establishing intestinal immune tolerance, we hypothesized that the presence of TCs might determine this developmental window. Our analysis of TC abundance in mice ranging from 7 days to 6 weeks of age revealed their striking enrichment between 1 and 3 weeks of age, with rapid decline thereafter (**Fig. 5a, Extended data Fig. 9a**). Notably, TCs, in particular TC IV, were enriched within the mLN compared to skin-draining peripheral LN (pLN) (**Fig. 5b**). To determine the dynamics of neonatal TC differentiation, we used *Rorc^Venus-creERT2^R26^lsl-tdtomato^* mice to label RORγt expressing cells and their progeny. Following treatment of mice with 4-OHT at P1, over 60% of TCs remained tdTomato^+^ at P7 (**Fig. 5c**). This proportion fell to 15% by P14, although total numbers of both tdtTomato^−^ and tdTomato^+^ TCs increased between P7–14, reflecting *de novo* TC differentiation or proliferation during this critical developmental window. Both the proportion of tdTomato^+^ TCs and total cell numbers declined from P14 reflecting waning differentiation beyond this age (**Fig. 5c, Extended data Fig. 9b)**. In contrast, the proportion of fate-mapped MHCII^+^ ILC3s declined between P7 and 14 but remained stable thereafter (**Extended data Fig. 9b**), consistent with the notion that ILC3s are maintained by self-renewal^39^. Thus, TCs or putative RORγt^+^ TC progenitors, are present at birth, and are prominently enriched within the mLN at the time of intestinal tolerance induction during early life. Together these data suggest a critical window of opportunity for pTreg cell differentiation, determined by a wave of TCs within mLN in early life.

**Figure 5.**
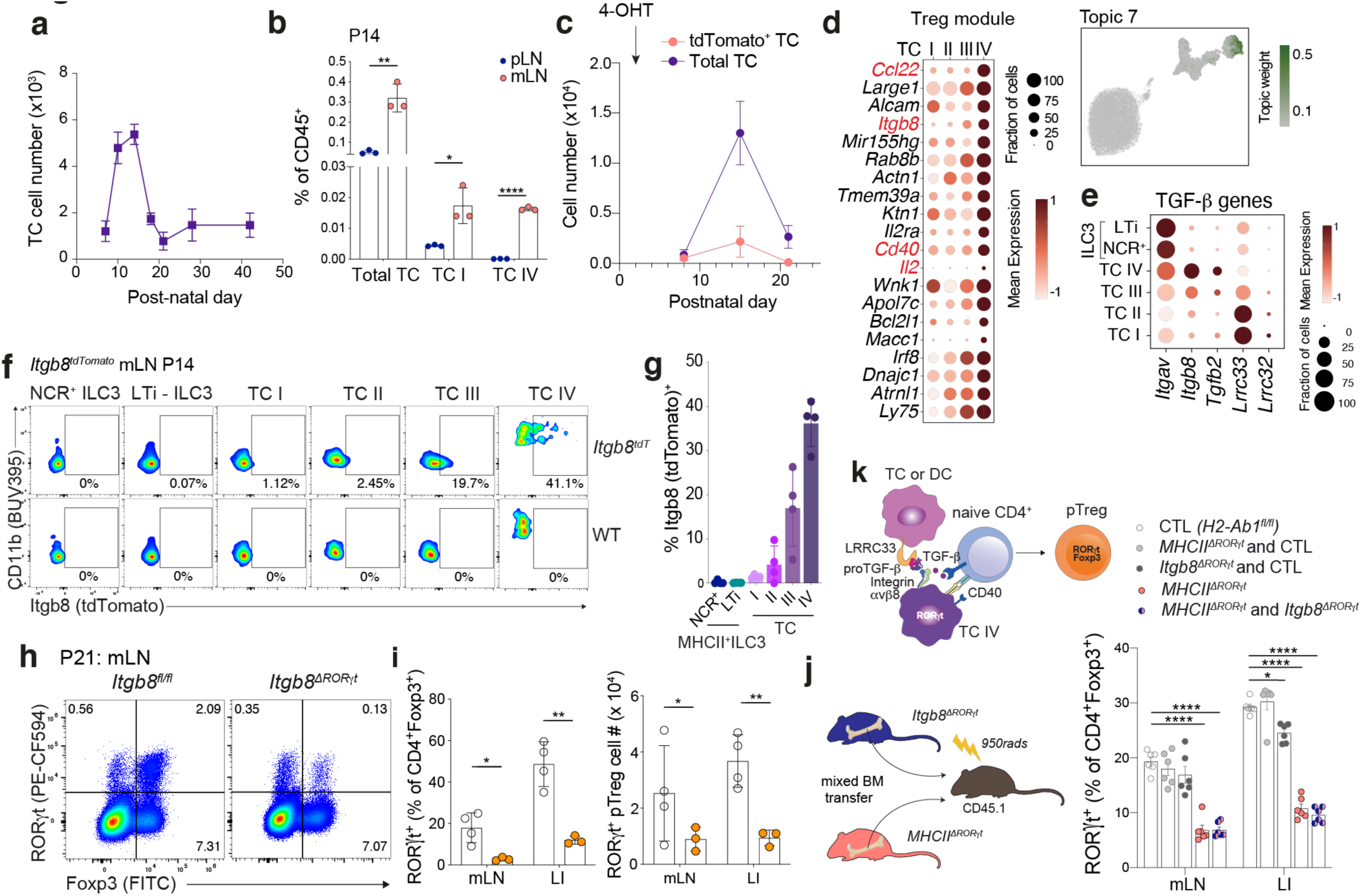
A developmental wave of TCs promotes early life pTreg differentiation in an Itgb8 dependent manner. **a**, Number of TCs within mLN from postnatal day 7 to 6-weeks-of age. Data pooled from two experiments with 3–8 individual mice per timepoint. **b**, Frequency of TCs in skin draining lymph nodes (pLN) and mLN of *Rorc^Venus-creERT2^*Aire^GFP^ mice at P14; *n* = 3 mice per group. **c**, Total number of tdTomato^−^ and tdTomato^+^ TCs isolated from mLN of *Rorc^Venus-creERT2^R26^lsl-tdTomato^*Aire^GFP^ mice at indicated time intervals following administration of 4-OHT on P1, *n* = 4–5 mice per timepoint. **d**, Topic modeling of 10X scRNA-seq TC transcriptomes. UMAP colored by the weight of topic 10 in each cell. **e**, Dot plot showing expression of TGF-β pathway genes in TCs and ILC3s. **f-g**, Representative flow cytometry (f) and summary graphs (g) of Itgb8 (Tdtomato) expression in TC and ILC3 subsets in mLN of *Itgb8^tdTomato^* (*n* = 4) or littermate wild-type (WT) mice. **h-i,** Representative flow cytometry of RORγt and Foxp3 expressing T cell subsets (**h**) and summary graphs for frequencies and numbers (**i**) of pTreg (RORγt^+^Foxp3^+^) cells in mLN and large intestine lamina propria (LI) of 3 w old *Itgb8^ΔRORγt^* (*n* = 3) and *Itgb8^fl/fl^* (*n* = 4) mice. **j**, Frequency of RORγt^+^ pTreg cells amongst CD4^+^Foxp3^+^ cells in mLN and LI of mixed bone marrow chimeras, analyzed 6 weeks post-reconstitution. *n* = 6 mice per group from 2 independent experiments. **k,** schema for pTreg induction by TCs. Error bars: means ± s.e.m. \**P* < 0.05; \*\**P* < 0.01; \*\*\*\**P* < 0.0001; unpaired two-sided *t*-test.

To determine if TC counterparts exist in humans, we analyzed a recent single-cell atlas encompassing second trimester to adult intestine and mLN^40^. Within a group of myeloid cells annotated as ‘lymphoid’ DCs we identified a cluster of cells distinct but closely related to CCR7^+^ DCs, expressing signature TC genes (*TNFRSF11B, SPIB*) including high levels of *AIRE* (**Extended data Fig. 9c-e**). Analysis of orthologous signature TC subset genes confirmed enrichment within the putative human TC cluster, most prominently for TC III and TC IV defining genes (**Extended data Fig. 9f**). In contrast to CCR7^+^ DCs, human TCs were almost exclusively present within the mLN (**Extended data Fig. 9g**) and were highly enriched within fetal samples (32% vs 3.8%; **Extended data Fig. 9h**), further implying a conserved role for TCs in early life intestinal tolerance.

The close temporal and spatial relationship between TC IV and pTreg cells supported a key role for this TC subset in pTreg differentiation. To determine if the TC IV subset has distinct features supporting pTreg generation, besides MHCII-dependent TCR stimulation, we used Latent Dirichlet Allocation (LDA), a probabilistic topic model, to capture shared and unique gene expression programs. Notably, the TC IV subset was enriched for a ‘Treg’ module encompassing critical molecules including IL-2, the TGF-β activating integrin Itgβ8, CD40 and Ccl22 (**Fig. 5d**). TGF-β signaling is known to be critical for pTreg cell differentiation. Activation of latent extracellular TGF-β requires physical interaction with either integrin αvβ6 or αvβ8 and loss of either TGF-β signaling on T cells, or Itgβ8 expression by hematopoietic cells, leads to impaired pTreg differentiation and development of autoimmunity and colitis^41,42^. Analysis of TGF-β signaling pathway genes in TC transcriptomes confirmed high expression of both *Itgav* and *Itgb8* by TC IV as well as unique expression of *Tgfb2* (**Fig. 5e**). To address the role of TCs in TGF-β mediated pTreg differentiation, we generated mice with conditional loss of the *Itgb8*gene in RORγt^+^ cells (*Itgb8^ΔRORγt^*). Importantly, although ILC3s expressed Itgav, they did not express Itgb8, as determined by analysis of SS2 single cell transcriptomes as well as *Itgb8^tdTomato^* reporter mice (**Fig. 5f,g, Extended data Fig. 10a,b**). Furthermore, ATAC-seq analysis showed that the *Itgb8* locus was inaccessible in ILC3s (**Extended data Fig. 10c**), confirming that *Itgb8^ΔRORγt^* mice have a specific deficiency of Itgb8 in TCs and T cells. Analysis of mLN and large intestine from 3-week-old mice, revealed a dramatic reduction in pTreg cell frequency and numbers (**Fig. 5h-i**), mirroring the loss of pTreg cells observed in *MHCII^ΔRORγt^* mice. Differentiation of pTreg cells was normal in *Cd4^cre^Itgb8^fl/fl^* mice (**Extended data Fig. 10d**), indicating that TGF-β activation by TCs, not T cells, is a critical mechanism for intestinal pTreg cell differentiation.

The unique expression of Itgb8 by TCs allowed us to test the requirement for TGF-β activation and antigen presentation by the same cell. Using bone marrow chimeras generated with a mix of *Itgb8^ΔRORγt^* and *MHCII^ΔRORγt^* bone marrow cells, in which both ILC3s and TCs can present antigen, but the same TC cannot present antigen and activate TGF-β, we found that pTreg differentiation was critically dependent on antigen presentation by Itgb8-expressing TCs, with an equivalent deficit in pTreg cells observed in *MHCII^ΔRORγt^* and *MHCII^ΔRORγt^/Itgb8^ΔRORγt^* chimeras (**Fig. 5j**), excluding the possibility of redundant and compensatory functions between TCs and ILC3s. Overall, our findings identify a new class of APCs, prominently present in the mLN during a critical early life window, and demonstrate an essential role for the TC IV subset in establishing intestinal tolerance through the generation of pTreg cells (**Fig. 5k**).

## Discussion

Contrary to the view that ‘neonatal immune privilege’, first demonstrated by Medawar in the 1950s^43,44^, results from the presence of immunosuppressive or ‘immature’ DCs with an inferior stimulatory capacity, our results suggest the existence of a dedicated tolerogenic APC type, enriched early in life. The requirement for MHC class II antigen presentation by TCs in early life, but not adulthood, provides support for a model in which a tightly regulated wave of TC differentiation during a narrow window in early life imprints durable microbiota-specific T cell tolerance. Intriguingly, pTreg cell abundance in adulthood is determined by cues sensed within the first week of life^45^, coincident with the observed wave of TC differentiation, suggesting that modulation of TC development may have lasting effects on intestinal immune tolerance.

A defining feature of TCs was their expression of RORγt. Our finding of overlapping markers between TCs and DCs resolves previous conflicting reports on RORγt^+^ DCs and their relationship to ILC3s^14,36^. Importantly, selective targeting of cDCs or ILC3s, enabled by identification of cell-type specific genes, demonstrated that neither ILC3s nor DCs, contribute to mucosal tolerance. While the precise ontogeny of TCs remains to be established, a notable finding was their expression of transcription factors and markers typically associated with mTECs. Intriguingly, a recent study highlighted the existence of hybrid cell types that emerge from Aire^+^ mTECs in the thymus^32^. Our discovery of TCs further challenges the current view of boundaries between cell lineages highlighting shared transcriptional programs between hematopoietic and non-hematopoietic cells that may support common purposes. Within the thymus, Aire^+^ mTECs instruct T cell tolerance through negative selection and neonatal thymic Treg generation^1,22^. The essential role of TCs in neonatal pTreg differentiation highlights symmetry between thymic and peripheral tolerance pathways. Given the expression of Aire by TC I and III, one can speculate that TCs may share additional functions with mTECs, such as tolerance to self-antigens.

In closing, our studies identified a novel tolerogenic APC type, enriched in the intestine during a critical early life period when host-microbiota symbiosis is first established. The finding that TC IV instructs extrathymic Treg cell generation provides a cellular basis for the reported early life window for establishment of intestinal immune tolerance. Future exploration of TC biology may yield key insights into mechanisms of immune tolerance and autoimmune and inflammatory disease pathogenesis.

## Acknowledgements

We thank Joris van der Veeken, Clarissa Campbell and members of the Rudensky lab for their technical support and helpful discussions. We thank the Single Cell Core Facility at MSK for sample processing. We acknowledge the use of the Integrated Genomics Operation Core, funded by the NCI Cancer Center Support Grant (CCSG, P30 CA08748), Cycle for Survival, and the Marie-Josée and Henry R. Kravis Center for Molecular Oncology. We thank Jay Gardner for discussions, Mark Anderson for provision of Aire^GFP^ mice, Caetano Reis e Sousa for *Clec9a^cre^* mice, Dean Sheppard for *Itgb8^fl/fl^* mice, Lisa Denzin for *H2-Dma^−/−^* mice, Michael Glickman for C7 mice and Sarah Teichmann for assistance with human single cell datasets.

## Author Contributions

C.C.B. and A.Y.R designed experiments and wrote the manuscript; Z.T. designed and performed computational analyses; B.A., G.S., Y.A.P.I, Y.F.P., L.F., V.T. and C.C.B. performed experiments and analyzed data; D.D. performed immunofluorescence staining and imaging analyses; S.H. and C.F. performed experiments; R.E. performed initial analysis of human intestinal single-cell data; H.A.P. performed electron microscopy analyses; J.V. processed tissues; M.K. performed immunofluorescence staining and imaging analyses for human tissue; L.J. generated mice under the supervision of M.vdB.; G.G. provided mice, J.C.M. provided mice and supervised experiments, T.G.P.G. supervised human IF experiments, C.L. supervised computational analyses and C.C.B. supervised experiments. All authors read and approved the manuscript.

## Materials and Methods

### Mice

*Rorc^Venus-T2A-creERT2^* mice were generated by insertion of a targeting construct into the *Rorc* 3-UTR by homologous recombination in embryonic stem (ES) cells on the C57Bl/6 background. The IRES-Venus-T2A-creER-frt-NeoR-frt cassette targeting construct was created by cloning. Homologous arms were retrieved from BAC clone RP24-209K20. To facilitate ES cell targeting Crispr/cas9 system was used. The gRNA was in vitro transcribed using MEGA shortscript T7 kit (Life Tech Corp AM1354) using recombineering techniques. The targeting vector, cas9 protein (Fisher Scientific A36498 Truecut Cas9 Protein v2) and gRNA were co-electroporated into G1 ES cells derived from an F1 hybrid blastocyst of 129S6 x C57BL/6J. The resulting chimeras were bred with FLPeR mice to excise the NEO cassette. *Rag1^RFP-creERT2^* (C57BL/6-Tg(Rag1-RFP,-cre/ERT2)33Narl) mice, obtained from the Rodent Model Resource Center (RMRC), were generated by insertion of a BAC transgene comprising the Rag1 promoter and RFP-IRES-creERT2 into ES cells from C57Bl/6 mice.

Adig(Aire^GFP^), *Clec9a^cre^, Rora^cre^, Itgb8^fl/fl^, Cd4^cre^, H2-Dma^−/−^,* C7 and *Itgb8^tdTomato^* mice have been previously described^24,46–52^. *Rorgt^cre^, H2-Ab1^fl/fl^, R26^lsl-tdTomato^, R26^lsl-YFP^*, *Zbtb46^GFP^, Il22^cre^,* C57Bl/6 (CD45.2), CD45.1 and BALB/c mice were purchased from Jackson Laboratories. Generation and treatments of mice were performed under protocol 21-05-007 and 08-10-023, approved by the Sloan Kettering Institute (SKI) Institutional Animal Care and Use Committee. All mouse strains were maintained in the SKI animal facility in specific pathogen free (SPF) conditions in accordance with institutional guidelines and ethical regulations. Both male and female mice were included in the study and we did not observe sex-dependent effects. All mice analyzed were age and litter matched unless otherwise specified. All animals used in this study had no previous history of experimentation and were naïve at the time of analysis.

### Tamoxifen diet

*Rorc^Venus-creERT2^H2-Ab1^fl/fl^* and littermate *Rorc^Venus-creERT2^H2-Ab1^fl/fwt^* mice were placed on a tamoxifen citrate–containing diet (TD.130860; Envigo) at 8 weeks of age for 5 weeks.

### Tissue processing

Mice were euthanized by CO_2_ inhalation. Organs were harvested and processed as follows. Lymphoid organs were digested in collagenase in RPMI1640 supplemented with 5% fetal calf serum, 1% L-glutamine, 1% penicillin–streptomycin, 10 mM HEPES, 1 mg/ml collagenase A (Sigma, 11088793001) and 1U/mL DNase I (Sigma, 10104159001) for 45 min at 37°C, 250 rpm. Large intestine was removed, flushed with PBS and incubated in PBS supplemented with 5% fetal calf serum, 1% L-glutamine, 1% penicillin–streptomycin, 10 mM HEPES, 1 mM dithiothreitol, and 1 mM EDTA for 15 min to remove the epithelial layer. Samples were washed and incubated in digest solution for 30 min. 1/4 inch ceramic beads (MP Biomedicals, 116540034) were added to large intestine samples (3 per sample) to aid in tissue dissociation. Digested samples were filtered through 100-μm strainers and centrifuged to remove collagenase solution. Thymus samples were minced with scissors followed by enzymatic digestion in RPMI1640 supplemented with 10% fetal calf serum, 1% L-glutamine, 10mM HEPES, 62.5ug/ml Liberase^TM^ and DNase I 0.4mg/ml. Density-gradient centrifugation using a three-layer Percoll gradient with specific gravities of 1.115, 1.065 and 1.0 was used to enrich for stromal cells for flow cytometric analysis. For sorting of mTECs, single cell suspension of digested thymocytes were depleted of CD45^+^ cells using CD45 microbeads (Miltenyi Biotec).

### Flow cytometry

For flow cytometric analysis, dead cells were excluded either by staining with LIVE/DEAD Fixable Violet or Zombie NIR in PBS for 10 minutes at 4°C, prior to cell-surface staining. Cells were then incubated with anti-CD16/32 in staining buffer (2% FBS, 0.1% Na azide, in PBS) for 10 minutes at 4°C to block binding to Fc receptors. Extracellular antigens were stained for 20-30 minutes at 4°C or RT (CCR7 staining) in staining buffer. For intracellular protein analysis, cells were fixed and permeabilized with Cytofix (BD Biosciences) and/or Ebioscience Foxp3 kit, per manufacturer instructions. Intracellular antigens were stained for 30min at 4°C in the respective 1x Perm/Wash buffer. Live cells were treated with DNase (0.08 U/ml) for 10min at RT and washed with staining buffer prior to acquisition on a BD LSR or Cytek Aurora™. 123count eBeads™ were added to quantify absolute cell numbers. The antibodies used for flow cytometry and FACS are listed in **Supplementary Table 4**. Unless otherwise stated, we used the following gatings: TCs: Lin (Siglec-F, TCRβ, TCRγ*δ*, CD19, B220, NK1.1)^−^CD64^−^Ly6C^−^RORγt (intracellular staining or expression of Venus in *Rorc^Venus-creERT2^ mice*)CXCR6^−^MHCII^+^; MHCII^+^ ILC3s: Lin^−^CD64^−^Ly6C^−^RORγt (intracellular staining or expression of Venus in *Rorc^Venus-creERT2^ mice*) CXCR6^+^MHCII^+^, and DC2s: Lin^−^ CD64^−^Ly6C^−^RORγt^−^CD11c^+^MHCII^+^CD11b^+^.

### Histological analysis of intestinal inflammation

Mice were euthanized by CO_2_ inhalation and large intestines were harvested and immediately placed into 10% formalin. Histopathological assessment for inflammation scoring in the intestine was performed on H&E stained sections based on established scoring systems for intestinal inflammation in mouse models^53^. Assessment includes severity and extent of inflammatory cell infiltrates, epithelial changes and mucosal architecture changes. Briefly, severity and extent of inflammatory cell infiltrate in the mucosa and if extending to submucosa and muscularis were evaluated histologically. Other evaluations include proliferation of epithelial cells lining the mucosa villous atrophy, crypts, loss of goblet cells, crypt abscesses, erosions and ulceration.

### Multiome scRNA and scATAC-sequencing

For single cell RNA and ATAC seq of RORγt^+^MHCII^+^ cells, mLN from 2 w old (P14–17) *Rorc^Venus-creERT2^* mice were pooled from 16 biological replicates and processed as described earlier. Cells were depleted of Lineage (TCRb, TCRγ*δ*, CD19, B220, NK1.1)^+^ cells via staining with biotinylated antibodies followed by magnetic bead negative selection. Cells were incubated with anti-CD16/32 in sorting buffer (2% FBS in PBS) for 10 minutes at 4°C to block binding to Fc receptors. Extracellular antigens were stained for 30 minutes at 4°C in sorting buffer (2% FBS, 2mM EDTA, in PBS). Cells were washed and resuspended in sorting buffer with SYTOX blue (Invitrogen) for exclusion of dead cells. Live, CD45^+^Lin(TCRb, TCRγ*d* CD19)^−^RORγt(Venus)^+^ MHCII^+^ cells were then sort purified. Cells were sorted into cRPMI, before being pelleted and resuspended in RPMI-2% FBS. Single Cell Multiome ATAC + Gene Expression was performed with 10X genomics system using Chromium Next GEM Single Cell Multiome Reagent Kit A (catalog no. 1000282) and ATAC Kit A (catalog no. 1000280) following Chromium Next GEM Single Cell Multiome ATAC + Gene Expression Reagent Kits User Guide and demonstrated protocol - Nuclei Isolation for Single Cell Multiome ATAC + Gene Expression Sequencing. Briefly, >50,000 cells (viability 95%) were lysed for 4min and resuspended in Diluted Nuclei Buffer (10x Genomics, PN-2000207). Lysis efficiency and nuclei concentration was evaluated on Countess II automatic cell counter by trypan blue staining. 9,660 nuclei were loaded per transposition reaction, targeting recovery of 6,000 nuclei after encapsulation. After the transposition reaction, nuclei were encapsulated and barcoded. Next-generation sequencing libraries were constructed following User Guide, which were sequenced on an Illumina NovaSeq 6000 system.

### Plate-based Smart-seq2 sequencing

RORγt^+^MHCII^+^ cells were enriched from a pool of MLN from 3-week-old (P21) *Rorc^Venus-creERT2^* mice. Cells were depleted of Lineage (TCRβ, TCRγ*δ*, CD19, B220, NK1.1)^+^ cells via staining with biotinylated antibodies followed by magnetic bead negative selection. Live, Lin(CD3, TCRβ, TCRγ*d* CD19, B220, NK1.1)^−^CD64^−^Ly6C^−^MHCII^+^RORγt(Venus)^+^ cells were then sorted into single wells. Cells were also stained for CD90, CD11c and CD11b for acquiring index sorting information on cell surface expression. Aire^+^ mTECs were enriched from a pool of thymi from 3-week old mice via staining with biotinylated antibodies against CD45 followed by magnetic bead negative selection. CD45^−^Epcam^+^MHCII^+^Aire(GFP)^+^ cells were sorted into single wells. Aire^+^ DCs were enriched from a pool of MLN from the same 3-week old mice. Cells were depleted of Lin^+^ cells as described above and live, Lin(CD3, TCRβ, TCRγ*d* CD19, B220, NK1.1)^−^CD90^−^CD64^−^Ly6C^−^CD11c^+^MHCII^+^Aire(GFP)^+^ cells were then sorted into single wells. Retrospective index sorting analysis confirmed that Aire(GFP)^+^ cells were CD11c^lo^MHCII^hi^, representing CCR7^+^ DCs.

Single cells were sorted into Buffer RLT (Qiagen). Cell lysates were immediately sealed and spun down before transferring to dry ice and storing at -80 °C. RNA was purified using the Agencourt RNAClean XP beads (Beckman Coulter) at a 2.2X ratio. First-strand cDNA synthesis was achieved using Maxima H Minus Reverse Transcriptase (ThermoFisher) according to the manufacturer’s protocol using oligo dT primers, with the addition of a custom template-switch oligo in a 1mM final concentration. cDNA was amplified for 24 cycles using KAPA HiFi HotStart ReadyMix (Kapa Biosystems KK2601). After PicoGreen quantification, 0.1-0.2ng of cDNA was used to prepare libraries with the Nextera XT DNA Library Preparation Kit (Illumina) in a total volume of 6.25µL with 12 cycles of PCR. Indexed libraries were pooled by volume and cleaned by aMPure XP beads (Beckman Coulter) at a 1X ratio. Pools were sequenced on a HiSeq 4000 in a PE50 or PE100 run using the HiSeq 3000/4000 SBS Kit (Illumina). An average of 1.8 million paired reads were generated per sample and the percent of mRNA bases per sample averaged 63%.

## Mouse single-cell RNA-seq and single-cell ATAC-seq computational analysis

### Pre-processing of the 10X multiome scRNA-seq and scATAC-seq for RORγt^+^MHCII^+^ cells

Single-cell RNA-seq and ATAC-seq FASTQ files were aligned to mm10 (Cell Ranger mouse reference genome mm10-2020-A-2.0.0) and counted by Cell Ranger ARC v2.0.0 with default parameters. The barcodes were filtered based on the number of RNA-seq transcripts (>1k and <50k), the number of detected genes (>500 and <6k), and the fraction of mitochondrial transcripts (<15%). Barcodes were further filtered based on the number of ATAC-seq fragments (3.5 < log10{nFrags} < 4.5) and TSS enrichment score (>4). Arrow files were created from the scATAC-seq fragments using ArchR v1.0.1^54^, and doublets were identified and removed with default parameters. Finally, any genes detected in <2 cells in the scRNA-seq data were discarded, leaving 20779 genes. After clustering the scRNA-seq data (described in ‘Dimensionality reduction, cell clustering, and visualization’), and based on the expression of marker genes, we identified 5 minor *Rorc*^−^ or *Ptprc^−^* contaminant clusters (glial cells; cluster 17, pDCs; cluster 18, Rorc^−^DCs; cluster 19, and mono/macrophages; clusters 20-21) which were excluded from downstream analyses. In total, 10145 cells remained, with a median scRNA-seq library-size of 3150 and a median of 13885 in the number of ATAC-seq fragments.

### Pre-processing of the Smart-seq2 scRNA-seq dataset

Smart-seq2 sequencing data from demultiplexed samples was aligned to the mouse reference genome using STAR v2.7.7a^55^ with ‘--twopassMode Basic --outFilterMultimapNmax 1 --quantMode TranscriptomeSAM’. Sequence reads were aligned and annotated using a STAR index created from GENCODE GRCm38 (mm10) release M25 primary assembly genome and gene annotations^56^. Alignment files were individually name-sorted using Samtools v1.11^57^, and then used to create a cell-by-gene count matrix using featureCounts^58^ (subread v2.0.1). The count matrix was filtered based on the number of transcripts (>50K), number of detected genes (>1300), and the fraction of mitochondrial transcripts (<8%). Finally, genes detected in <2 cells were discarded. A total of 481 cells remained, with a median library size of 924319 from 27195 genes.

### Dimensionality reduction, cell clustering, and visualization

For each scRNA-seq dataset, the filtered count matrix was library-size normalized, log-transformed (‘log-normalized’ expression values) and then centered and scaled (‘scaled’ expression values) using Seurat v4.0.4. Principal component analysis (PCA) was performed on the scaled data (npcs=50). PhenoGraph clustering^59^ was performed using the first *N* principal components (PCs) with *k* nearest neighbors (*N*=30 and *k*=30 for the multiome scRNA-seq data; *N*=20 and *k*=30 for the Smart-seq2 dataset; *N*=30 and *k*=20 for the human gut DCs). Cell clustering was visualized using UMAP^60^, computed from the nearest neighbor graph built by PhenoGraph.

The multiome scATAC-seq data analysis was restricted to the cells in clusters 1-16 of the scRNA-seq results, as previously described for pre-processing. Latent Semantic Indexing (LSI) was performed on 100000 top variable tiles (500 bp genomic bins) identified after 10 iterations of ‘IterativeLSI’ by ArchR. Tiles from non-standard chromosomes, chrM, and chrY were not included in this analysis. Cells were clustered (method=Seurat, k.param=30, resolution=1.2) and visualized with UMAP (nNeighbors=30) using 30 LSI components. In both the scRNA-seq and scATAC-seq data, we identified several clusters of LTi cells (scRNA clusters 9-16 and scATAC clusters 7-13). These clusters showed weak pairwise matchings between scRNA and scATAC; therefore, they were combined as one group of LTi cells for downstream analyses, where stated.

### Differential gene expression tests

Differentially expressed genes (DEGs) between groups of cells were identified with MAST^61^, performed using Seurat functions. MAST was run on the log-normalized expression values. In all tests, genes were only considered if they were detected in at least 10% of the cells in at least one of the two groups compared (min.pct=0.1, logfc.threshold=0). In one-vs-rest DE tests comparing multiple groups, each group was compared to all the cells from other groups. Specific DE comparisons are described in the results. DEGs were reported according to their fold change (>1.5) and adjusted *p*-value (<0.01). Ribosomal and mitochondrial genes were removed from the final list of genes reported/visualized. Where stated, the top DEG markers were subsequently selected for each group, based on fold change.

### Data imputation for scRNA-seq data

MAGIC imputation^62^ was applied to the log-normalized expression values for the multiome scRNA-seq dataset to further de-noise and recover missing values. Imputed gene expression values were only used for data visualization on UMAP overlays and heatmaps, where stated.

### Cell-cycle scores

Using standard Seurat functions, we computed cell cycle scores for known S-phase and G2/M-phase marker genes^63^ to identify proliferating cells.

### Topic modeling for scRNA-seq data

‘Topics’ were identified by fitting a Latent Dirichlet Allocation (LDA) model, also known as a Grade of Membership (GoM) model^64^, to the raw gene expression count matrix for TCs (clusters 1-5 of the multiome scRNA-seq data) using CountClust v1.18.0^65^. Genes that were detected in fewer than 10 TCs were not included. The optimal number of topics (*K*=8) was selected among values ranging from 3 to 15 with the maximum Bayes Factor (BF). The role of a topic in each cell is measured by the degree to which it represents that topic, and the topic weights sum up to 1 in each cell. The importance of a gene for each topic is measured by how distinctively differentially expressed it is in that topic, by measuring the KL-divergence of its relative gene expression to other topics, assuming a Poisson distribution. One topic, defined by ribosomal and mitochondrial genes and shared across all clusters, was removed from the topic model visualizations.

### Dynamical modeling of RNA velocity for the multiome scRNA-seq data

The unspliced and spliced mRNAs for the scRNA-seq profiles of the multiome data were counted by Velocyto v0.17.17^30^ from the position-sorted BAM file containing GEX read alignments, outputted by Cell Ranger ARC in pre-processing. As annotation files for Velocyto, we used the same mm10 gene annotations used in pre-processing, in addition to the mm10 expressed repeat annotation from the “RepeatMasker” track of UCSC genome browser. Next, we used the Velocyto results to learn a generalized dynamical model of RNA velocities by scVelo v0.2.4^66^. Count matrices were filtered, normalized, and log transformed (min_shared_counts = 10, n_top_genes = 3000), cell-cycle effect was corrected by regressing out S-phase and G2/M-phase scores, using Scanpy 1.6.0^67^. After performing PCA on the corrected data (n_pcs = 30), first- and second-order moments were computed for each cell across its nearest neighbors in the PCA space (n_neighbors = 30). Finally, the full splicing kinetics were recovered and solved for each gene by scVelo’s dynamical model.

### Integrating the Smart-seq2 dataset with the multiome dataset

RORγt^+^MHCII^+^ transcriptomes (based on cell-type as sorted) from the SS2 dataset were integrated with transcriptomes from the 10X multiome scRNA-seq data, using Seurat^68^. Based on the variability of genes in both datasets, 5000 top scoring genes were selected by Seurat functions to identify ‘integration anchors’ with Canonical Correlation Analysis (CCA). Expression values for these genes were integrated, scaled, and used for PCA. A UMAP embedding was computed from the first *N*=30 PCs (*k*=30). Additionally, using Seurat functions, the RORγt^+^MHCII^+^ cells from the SS2 dataset (query) were mapped to multiome scRNA-seq clusters (reference) by projecting the PCA from the reference onto the query to identify ‘transfer anchors’, and then assigning a prediction score for each reference cluster to query cells. The cluster identity with the highest score is chosen as the predicted label for each cell.

### Single-cell enrichment scores for gene sets

Given a set of genes, we standardized the log-normalized expression values of each gene across cells and then averaged these values for all genes in the set, assigning an enrichment score to each cell. Where stated, these scores were standardized across cells and reported as *z*-scores.

### Creating pseudo-bulk samples from scRNA-seq data

Pseudo-bulk samples were created by averaging the unimputed log-normalized gene expression values for each cluster. In cases where scaled values were used for downstream analyses, these average expression values were standardized across the pseudo-bulk samples.

### Tissue-restricted antigen (TRA) enrichment in multiome scRNA-seq data

An established list of 6611 TRA genes was taken from previously published data^33^ and filtered to 4587 TRA genes detected in the multiome scRNA-seq dataset. We computed an enrichment score of this gene set for each cell, excluding proliferating cells (cluster 6 and 7) and ILC3p cells (cluster 1).

### Identifying TC-enriched TRAs by gene set enrichment analysis (GSEA)

To avoid potential noise from single-cell data in GSEA, we created pseudo-bulk samples for clusters in the multiome scRNA-seq data. GSEA v4.1.0^69,70^ was performed to compare the enrichment of TRAs in non-proliferating TC samples (clusters 2-5) versus LTi samples (clusters 9-16), with ‘log2_Ratio_of_Classes’ as the gene ranking metric and otherwise default settings. 1554 TRAs with ‘core enrichment’ were selected as top TRAs for TCs.

### Similarity of multiome scRNA-seq clusters to bulk microarray ImmGen samples

The RMA-normalized and log2-transformed gene expression data of 224 bulk microarray samples from a publicly available ImmGen dataset was downloaded from https://www.haemosphere.org^71^. For each gene, the probeset with the highest mean expression was retained. We included all cell types isolated from naïve, untreated mice. Pseudo-bulk samples were generated from the multiome scRNA-seq data for each TC subset, and non-proliferating MHCII^+^ ILC3s (NCR^+^ ILC3 and LTi cells). The gene expression vectors were scaled across bulk/pseudo-bulk samples within each dataset, and their pairwise cosine similarities were used to compare the samples. These similarity scores were computed from the expression of 2399 DEGs (FC >1.3, adjusted *P* <0.01) comparing the scRNA-seq clusters in a one-vs-rest test, that were also expressed in the microarray data. The proliferating and progenitor clusters were excluded from the DE test, and the LTis were grouped together.

### Cell Type Prediction

To determine similarity between TCs and known cell types we used CellTypist (https://www.celltypist.org), with both low- and high-resolution models of immune cells to classify cells with coarse and fine granularities, respectively. Top predicted labels for each input cell were visualized.

### Similarity of TCs to thymic epithelial cells

scRNA-seq profiles of CD45^−^ thymic epithelial cells were downloaded from a publicly available dataset (GSE103967)^20^. The raw counts were library-size normalized, log-transformed, and used to create pseudo-bulk samples for each thymic epithelial cluster. Pseudo-bulk samples were also generated to represent the multiome scRNA-seq TC clusters (2-5). These pseudo-bulk gene expression vectors were scaled across samples within each dataset, and their pairwise cosine similarities were used to compare clusters from the two datasets. These similarity scores were computed from the expression of 1740 DEGs (FC >1.3, adjusted *P* <0.01) identified in a one-vs-rest DE test for non-proliferating TC clusters (2-5), that were also expressed in the thymic epithelial cells. Amongst individual clusters of thymic epithelial cells defined in the original dataset, we identified 2 clusters of transit amplifying Aire^+^ cells (clusters 25 and 26), distinguished by signature gene expression including cell cycle genes.

### scRNA-seq dataset of human gut DCs

Dendritic cells (annotated as cDC1, cDC2, or Lymphoid DC) within the myeloid dataset from the human gut atlas^40^ were re-clustered. From the gene markers for each TC subset (one-vs-rest DE test for non proliferating TC scRNA-seq clusters, FC >1.5, adjusted *P* <0.01, we identified orthologous human genes that were uniquely mapped by gprofiler2 and computed enrichment scores for TC sub-set gene signatures for each human cell.

### Peak calling for the multiome scATAC-seq data

For peak-calling of the scATAC-seq data, clusters for similar cell types were grouped: C1 (TC IV), C2-4 (TC I,II,III), C5-6 (NCR+ ILC3), and C7-13 (LTi). Filtered ATAC-seq fragments for each group were extracted from ArchR arrow files. We performed MACS2 v2.2.7.1 on fragments of each group with ’-- gsize mm --qval 0.01 --nomodel --ext 200 --shift -100 --call-summits’. The peak summits were extended by 100 bp in each direction. Regions extending outside of mm10 chromosomes, arising from chrY or chrM, overlapping with blacklist regions precompiled by ArchR (merged from the ENCODE mm10 v2 blacklist regions from https://github.com/Boyle-Lab/Blacklist/blob/master/lists/mm10-blacklist.v2.bed.gz and mitochondrial regions that are highly mappable to the mm10 nuclear genome from https://github.com/caleblareau/mitoblacklist/blob/master/peaks/mm10_peaks.narrowPeak), or containing ‘N’ nucleotides (>0.001 of the sequence) were filtered. Regions from all groups were compiled and overlapping regions were merged to their union, resulting in a non-overlapping set of 176942 peaks. A peak-by-cell count matrix was created by ArchR with a ‘ceiling’ value of 4 for the counts to avoid strong biases.

### Transcription factor (TF) motif enrichment with chromVAR

The peaks that were accessible in <10 cells were filtered from the peak insertion counts, created as described in the previous section, and the resulting 176898 x 10145 peak-by-cell count matrix was used for motif enrichment with chromVAR v1.14.0^72^. Mouse motif PWMs were downloaded from the CIS-BP database^73^ (‘Mus_musculus_2022_01_14_6-40_pm’), and the missing PWMs were extracted from ‘mouse_pwms_v1’ in chromVARmotifs v0.2.0. The GC content of the peaks was computed with chromVAR, and motifs were matched to them by motifmatchr v1.14.0. Then, chromVAR ‘deviations’ of the motifs were computed for the peak-by-cell count matrix. The ‘top motif’ for each TF was selected by correlating its log-normalized gene expression values (from multiome scRNA-seq) with the deviation *z*-scores of its motifs, in the same cells, and picking the motif with the highest Pearson correlation coefficient. Finally, TF-motif pairs with a correlation higher than 0.1 were selected. This resulted in 56 top TFs, out of 739 CIS-BP TFs that were expressed (i.e. had any transcripts detected) in the multiome scRNA-seq profiles. The same process was repeated for the 139528 x 1552 peak-by-cell count matrix of TCs (multiome scATAC-seq clusters 1-4) and the peaks accessible in at least 10 TCs. Out of 652 CIS-BP TFs that were expressed in TCs, 68 had a TF-motif correlation higher than 0.1 and were selected as top TFs for TCs.

### Neonatal 4-OH Tamoxifen administration

For labeling of RORγt^+^ cells, *Rorc^Venus-creERT2^*Aire^GFP^ mice were injected intra-peritoneally (i.p.) on P1 with 25μg 4-OH-tamoxifen (4-OHT) and analyzed on P8, 15 and 21. For RAG1 fate-mapping, *Rag1^RFP-creERT2^R26^lsl-YFP^* mice were injected with 25μg 4-OHT i.p. on P3, 5 and 7 and analyzed on P15.

### Electron microscopy

RORγt^+^MHCII^+^ cells were enriched from a pool of mLN from P18 *Rorc^Venus-creERT2^* mice (for TC IV or reference CCR7^−^ and CCR7^+^ DCs) and P18 Aire^GFP^ mice (for TC I). Cells were depleted of Lineage (TCRb, TCRgd, CD19, B220, NK1.1)^+^ cells via staining with biotinylated antibodies followed by magnetic bead negative selection. mTECs were enriched from a pool of thymi from P18 AireGFP mice as described above. Live, Lin(TCRb, TCRgd, CD19, B220, NK1.1)^−^CD64^−^Ly6C^−^ MHCII^+^RORγt(Venus)^+^CD11c^+^CD11b^+^cells (TC IV), Lin^−^RORγt(Venus)^−^CD11c^lo^MHCII^hi^ (CCR7^+^ DC), Lin^−^RORγt(Venus)^−^CD11c^hi^MHCII^−^ (CCR7^−^ DC), Lin^−^CXCR6^−^CD11c^−/lo^Aire(GFP)^hi^ (TC I), Lin^−^ RORγt(Venus)^+^CD90^+^MHCII^+^ (LTi/ILC3), or CD45^−^Epcam^+^MHCII^+^Aire(GFP)^+^ cells were then sorted directly into 2% glutaraldehyde, 4% PFA, and 2 mM CaCl_2_ in 0.1 M sodium cacodylate buffer (pH 7.2), fixed for >1 h at room temperature, postfixed in 1% osmium tetroxide, dehydrated in acetone, and processed for Epon embedding. Ultrathin sections (60–65 nm) were counterstained with uranyl acetate and lead citrate. Images were taken with a transmission electron microscope (Tecnai G2-12; FEI, Hillsboro, OR) equipped with a digital camera (AMT BioSprint29).

### Immunofluorescence: tissue preparation, microscopy, and image analysis

Mesenteric lymph nodes (mLN) were dissected from 2–3 week-old *Rorc^Venus-creERT2^* or *AIRE^GFP^* mice and trimmed of fat using a dissection scope and forceps. mLNs were fixed in 2% paraformaldehyde for 4 hours in 4°C, washed 3x in PBS, and dehydrated in 30% sucrose in 0.1 M phosphate buffer overnight (16-20 hours). mLNs were embedded in optimal cutting temperature (OCT) compound, frozen on dry ice, and stored at -80°C. 15-20m sagittal sections were placed on Superfrost Plus microscopy slides and stored at -20C until staining. mLN sections were permeabilized using 0.2% Triton X-100 for 15 minutes at room temperature, washed 3x with PBS, blocked in 5% rabbit and donkey serum for 1 hour at room temperature, and washed 3x with PBS. Next, the sections were incubated with combinations of the following primary antibodies in PBS overnight at 4°C: CD11c BV421 (Biolegend, Clone N418, 1:50), CD11b BV480 (BD Biosciences, Clone M1/70 1:50) CD4 AF647 (Biolegend, Clone RM4-5, 1:50), Foxp3 eFluor 570 (ThermoFisher, Clone FJK-16s, 1:50), GFP AF488 (ThermoFisher A12311, polyclonal, 1:100), Rorgt APC (ThermoFisher, Clone AFKJS-9, 1:50), and MHCII AF700 (Biolegend, Clone M5/114.15.2, 1:200) antibodies. The samples were washed 3x in PBS the next day and mounted in SlowFade Diamond antifade reagent (ThermoFisher). No. 1.5 coverglass was used to seal the slide and all subsequent imaging was done on Leica SP8 microscope. Analysis was performed by histo-cytometry methods using Imaris software. Image segmentation was performed in Imaris using the “Surface Object Creation” module, which employs a seeded region growing, k-means, and watershed algorithm to define individual cells.

### In vitro cell culture

Naïve CD4^+^vβ10^+^CD25^−^CD44^lo^CD62L^hi^ C7 T cells were sort purified after enrichment with a CD4^+^ T-cell negative selection kit (Miltenyi Biotec). DC and TC subsets were sort purified from *Rorc^Venus-creERT2^* mice using gating strategy above with inclusion of CCR6 to distinguish TC I and II from TC III and IV. cDC2s were distinguished from their CCR7^+^ counterparts by CD11c^hi^MHCII^int^ expression. T cells were co-cultured in triplicate at a ratio of 300 DC or TCs to 1 × 10^3^ T cells in the presence of ESAT peptide (1µg/ml; Invivogen), with the addition of 0.5ng/ml TGF-β1 (Peprotech), 100IU/ml of IL-2 (NCI). Foxp3 expression was assessed after 4 days of culture.

### Bone marrow chimera mice

Bone marrow (BM) cells were isolated from indicated donor mice, depleted of CD90.2^+^ and TER-119^+^ cells using magnetic bead based depletion. BM cells were resuspended in PBS and 2-3 x 10^6^ cells were injected into 6-week-old CD45.1 mice, irradiated with 950rads/mouse one day earlier. After 6 weeks, mLN and LI LP were harvested for analysis.

### Statistical analysis

Analysis of all data was done with unpaired two-tailed t test, one or two-way ANOVA with a 95% confidence interval, or Mann-Whitney U test, as specified in the text or legends. *P* < 0.05 was considered significant: * *p* < 0.05; ** *p* < 0.01; \*\*\**p* < 0.001; \*\*\*\**p* < 0.0001. Details as to number of replicates, sample size, significance tests, and value and meaning of *n* for each experiment are included in the Methods or Figure legends. Statistical tests were performed with Prism (GraphPad Software). scATAC and RNA-sequencing experiments were carried out once. Unless otherwise stated, all other experiments were carried out independently at least twice. Mice were non-randomly allocated to experimental groups to ensure equal distribution of genotypes between treatments. Researchers were not blinded as to genotype or treatment during the experiments. No measures were taken to estimate sample size of to determine whether the data met the assumptions of the statistical approaches used. Significance (α) was defined as < 0.05 throughout, after correcting for multiple comparisons.

### Data and material availability

The mouse sequencing data are available through the Gene Expression Omnibus under accession GSExxxx.

## List of Supplementary Tables

**Supplementary Table 1**. Table_S1.csv

Excel file containing top 50 marker genes for mTECs vs TCs (SS2).

**Supplementary Table 2**. Table_S2.csv

Excel file containing top marker genes for SS2 clusters

**Supplementary Table 3.** Table_S3.csv

Excel file containing list of TC-enriched tissue restricted antigens.

**Supplementary Table 4**. Table_S4.xlsx

Excel file containing list of antibodies used in this study.

---

**Extended Data Figure 1.**
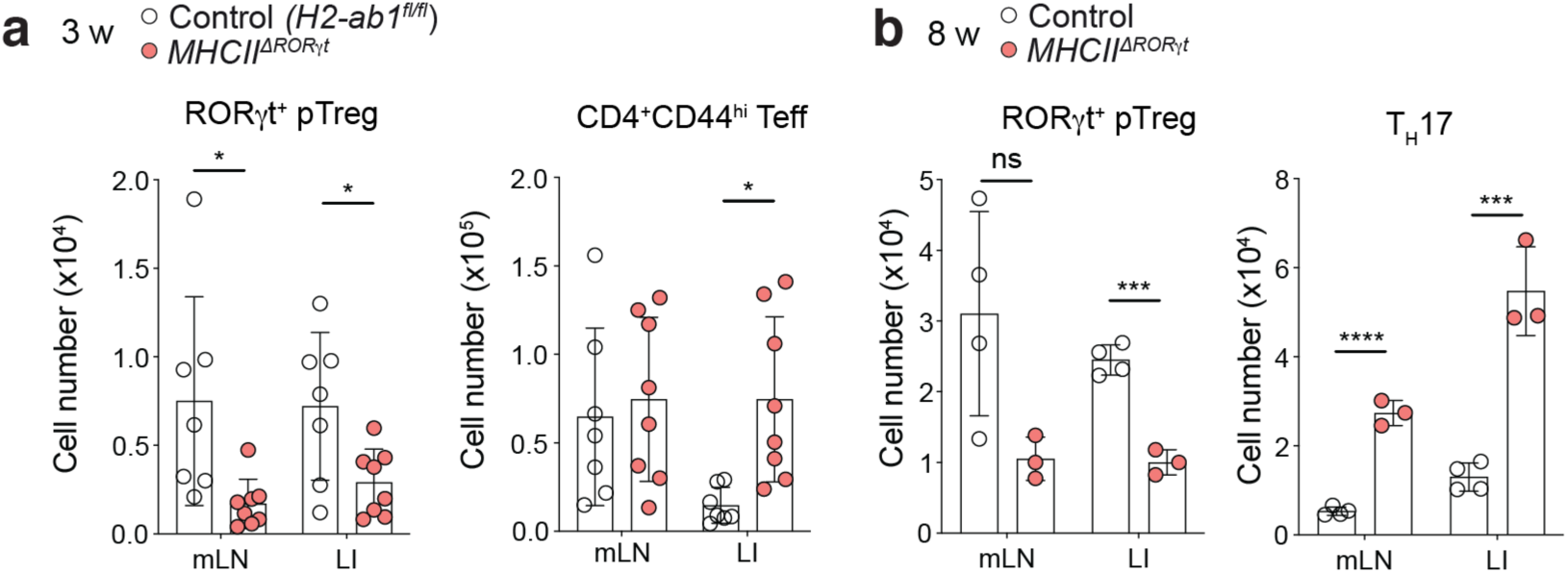
Analysis of pTreg cell generation in mice harboring MHC class II-deficient RORγt^+^ APCs. **a**, Quantification of total pTreg (RORγt^+^Foxp3^+^) and CD4^+^ T_eff_ (Foxp3^−^CD44^hi^) cells in the mesenteric lymph nodes (mLN) and large intestine lamina propria (LI) of 3-week-old *MHCII^ΔRORγt^* and control (*H2-Ab1^fl/fl^*) mice (*n* = 7 or 8 mice per group from two independent experiments). **b**, pTreg (RORγt^+^Foxp3^+^) and T_H_17 (Foxp3^−^CD44^hi^RORγt^+^) cells in mLN and LI of 8-week-old *MHCII^ΔRORγt^* (*n* = 4) and control (*n* = 3) mice. Error bars: means ± s.e.m.. \**P* < 0.05; \*\*\**P* < 0.01, \*\*\*\**P* < 0.0001; unpaired two-sided *t*-test.

**Extended Data Figure 2.**
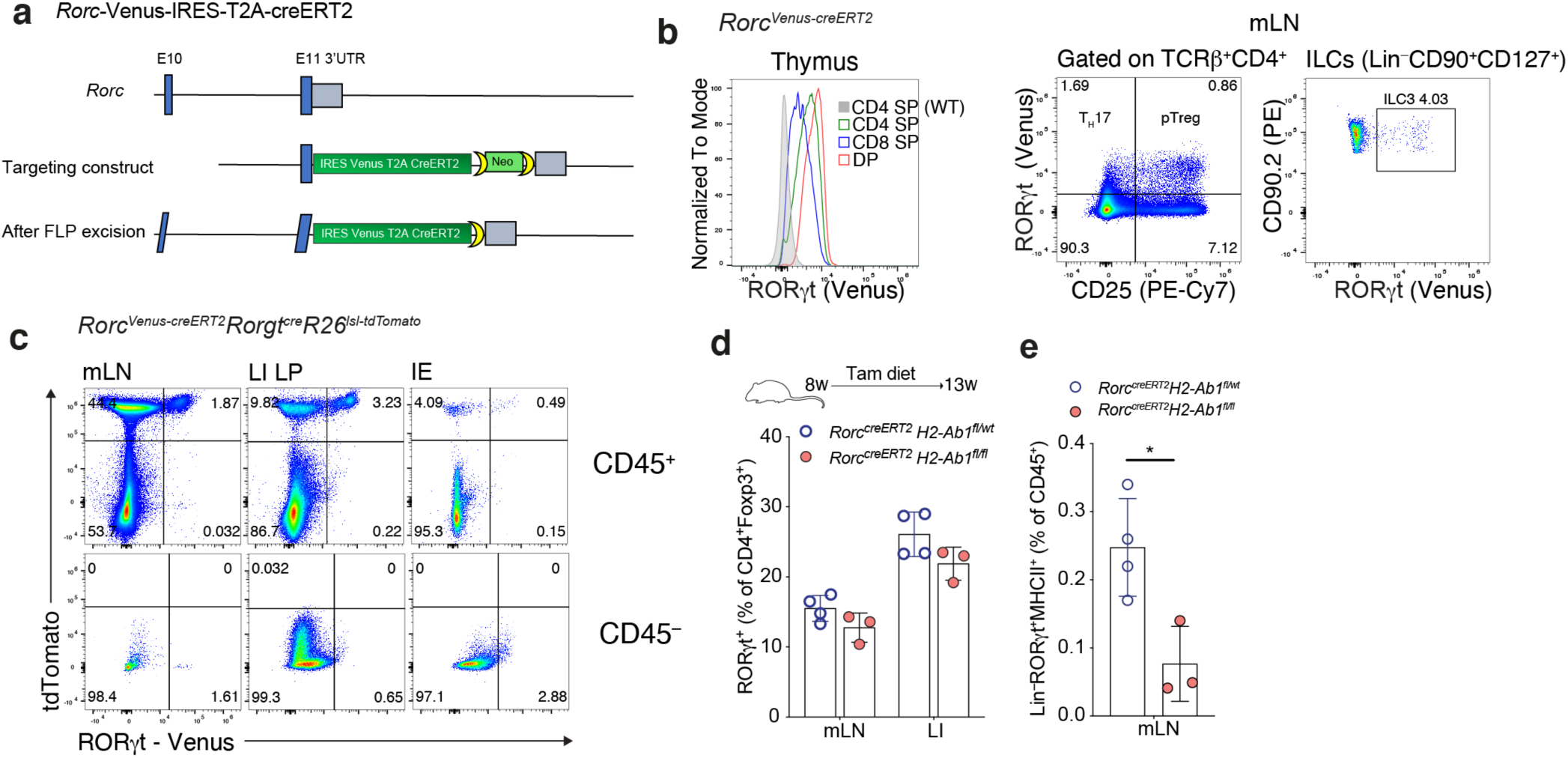
Temporal ablation of MHC Class II on RORγt^+^ APCs. **a,** Targeting strategy for the *Rorc* locus. **b,** Flow cytometry of Venus expression in thymocytes (left) or mLN TCRβ^+^CD4^+^ T cells (middle) and Lin^−^CD90^+^CD127^+^ innate lymphoid cells (ILC; right) isolated from adult mice. **c,** Flow cytometry of mLN, LI and IEL CD45^+^ and CD45^−^ cells in P16 RORγt fate-mapper Rorc reporter (*Rorc^Venus-creERT2^Rorgt^Cre^Rosa26^lsl-tdT^*) mice. Representative of six mice from two independent experiments. **d-e**, Frequency of pTreg cells amongst CD4^+^Foxp3^+^ cells in mLN and LI (d) or frequency of RORγt^+^ APCs (Lin–RORγt(Venus)^+^MHCII^+^) (e) in mLN of *Rorc^Venus-creERT2^H2-Ab1^fl/fl^* (n = 4) or *Rorc^Venus-creERT2^H2-Ab1^fl/wt^* (*n* = 3) mice maintained on tamoxifen diet from 8–13 weeks of age. **d**, Each symbol represents an individual mouse. Error bars: means ± s.e.m.; \**P* < 0.05; unpaired two-sided *t*-test.

**Extended Data Figure 3.**
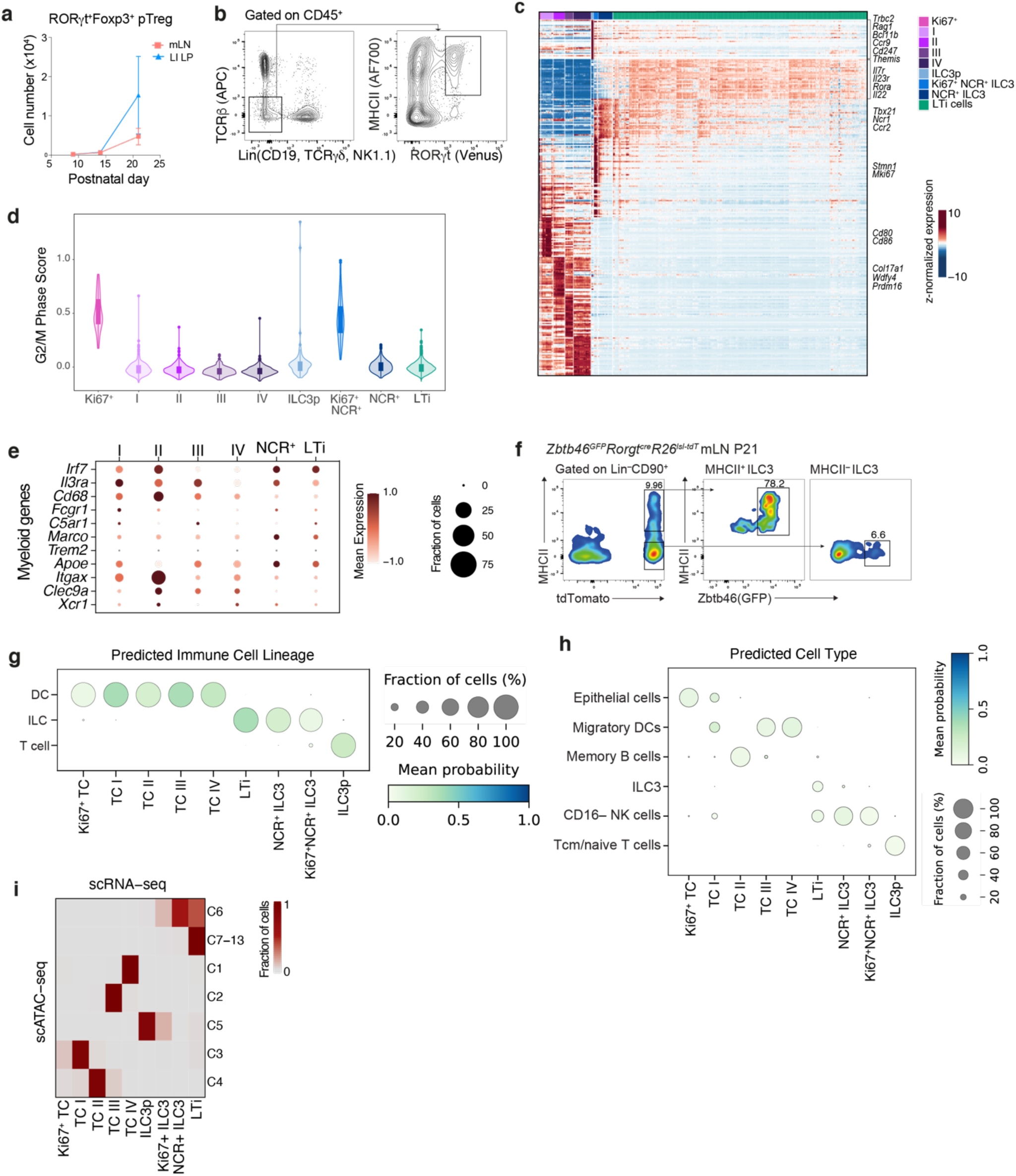
Identification of a novel RORγt^+^ APC lineage. **a**, RORγt^+^Foxp3^+^ pTreg cell numbers in mLN and LI of *Rorc^Venus-creERT2^* mice at indicated postnatal ages, *n* = 3–4 mice per time-point. **b**, Cell sorting scheme for Lin(TCRβ, TCRγ*δ*, CD19, NK1.1)^−^ RORγt(Venus)^+^MHCII^+^ cells. **c**, Heatmap reporting scaled, imputed expression of top differentially expressed genes for each scRNA-seq cluster (one vs the rest, FC>1.5, *P*<0.01). **d**, Expression score of cell-cycle genes for each scRNA-seq cluster. **e**, Dot plot showing expression of myeloid genes. **f**, Flow cytometric analysis of Zbtb46 (GFP) expression in ILC3 subsets from mLN of 3-week-old *Zbtb46*^GFP^*Rorgt^cre^R26^lsl-tdtomato^* mice. Representative of four mice from two independent experiments. **g-h**, CelltTypist derived cell labels for broad classification (g) and finer cell type annotation (h). **i**, Correspondence between cell labels for scATAC-seq and scRNA-seq.

**Extended Data Figure 4.**
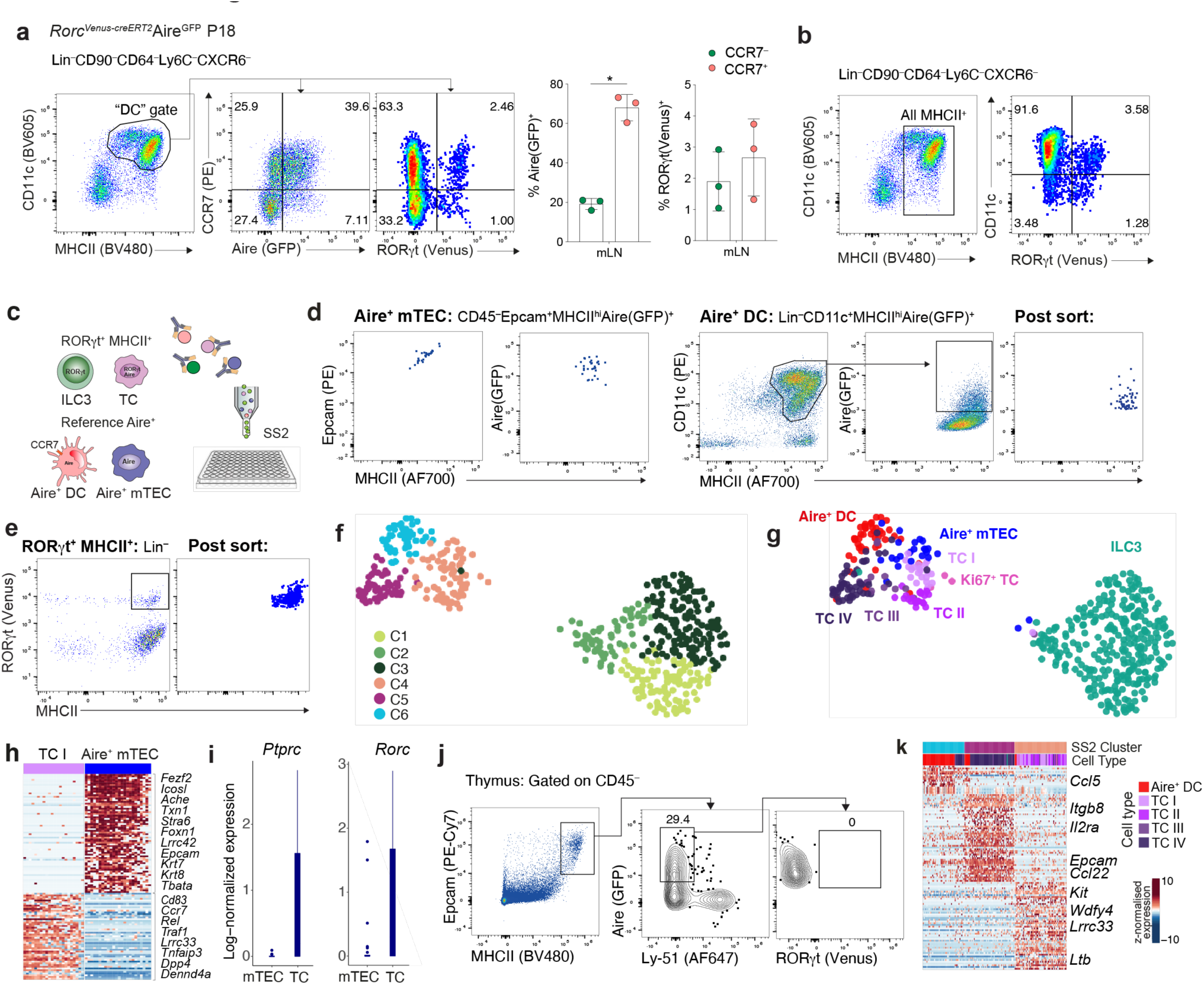
Distinguishing features of TCs, mTECs and DCs. **a**-**b**, Flow cytometric analysis of Lin^−^CD64^−^Ly6C^−^CD11c^+^MHCII^+^ ‘DCs’ (**a**) and Lin^−^ CD64^−^Ly6C^−^CXCR6^−^ RORγt(Venus)^+^MHCII^+^ TCs (**b**) in mLN of *Rorc^creERT2-Venus^*Aire^GFP^ mice at P18. **c,** Schematic of single cell transcriptome profiling of RORγt^+^MHCII^+^ cells from mLN of P21 *Rorc^Venus-creERT2^* mice encompassing TCs and MHCII^+^ ILC3s, alongside reference mLN Aire^+^MHCII^hi^ (CCR7^+^) DCs and thymic Aire^+^ mTECs from P21 Aire^GFP^ mice. **d**, Flow cytometric analysis of index sorted thymic Aire^+^ mTECs, mLN Aire^+^ DCs isolated from 3-week-old Aire^GFP^ mice and (**e**) mLN Lin^−^RORγt^+^MHCII^+^ cells from 3-week-old *Rorc^Venus-creERT2^* mice. **f**-**g**, UMAP visualization of 481 cells colored by (**f**) PhenoGraph cluster or (**g**) reference cell-type or RORγt^+^MHCII^+^ cell-type as assigned by mapping RORγt^+^MHCII^+^ SS2 cells to 10X scRNA-seq clusters (Fig. 2b). **h**, Heatmap reporting scaled expression values for top differentially expressed genes (FC>1.5, adj. *P*<0.01) between Aire^+^ mTEC and TC I. **i,** Bar graph showing log-normalized expression of *Ptprc* and *Rorc* genes in Aire^+^ mTECs and TCs. **j**, Flow cytometry of RORγt expression in Aire^+^ mTECs isolated from P18 *Rorc^Venus-creERT2^*Aire^GFP^ mice. Representative plot from one of two independent experiments with *n* = 3–5 mice. **k**, Heatmap reporting scaled expression values for top differentially expressed genes (FC>1.5, adj. *P*<0.01) between indicated SS2 clusters (f).

**Extended Data Figure 5.**
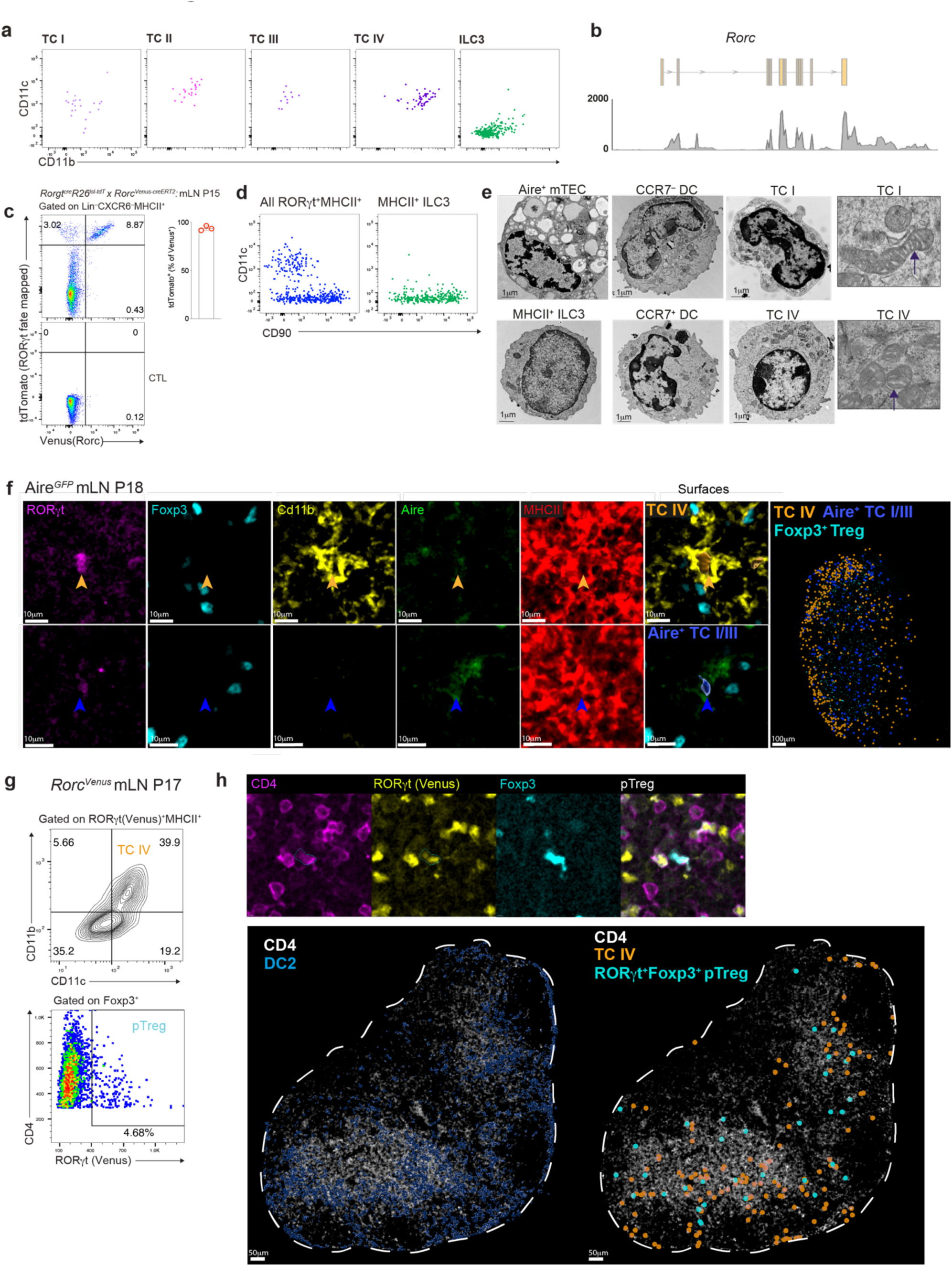
Phenotypic characterization of Thetis cells. **a**, Flow cytometry of index sorted mLN RORγt^+^MHCII^+^ cells for canonical DC markers, CD11c and CD11b**. b**, Coverage track for smart-seq2 single cell sequencing reads mapping to the *Rorc* locus, demonstrating expression of the Rorgt isoform by TCs. **c,** Flow cytometry of Lin^−^CXCR6^−^MHCII^+^ cells from mLN of *Rorgt^Cre^R26^lsl-tdtomato^Rorc^Venus-CreERT2^* mice and summary graph of frequency of tdTomato^+^ cells amongst Lin^−^CXCR6^−^Venus(YFP)^+^ cells (*n* = 3 mice)**. d**, Index sorting flow cytometric analysis of all RORγt^+^MHCII^+^ cells (left panel) and cells identified as ILC3 (right panel). **e**, Electron microscopy of CCR7^−^ DCs, CCR7^+^ DCs, Aire^+^ mTECs, TC I, TC IV and MHCII^+^ ILC3 cells. Far right panel: arrows indicate distinctive mitochondrial cristae in TCs. **f**, Representative immunofluorescence imaging of TC and Treg markers in mLN sections from 2-week-old Aire^GFP^ mice. Arrowheads indicate Aire^+^ TC I/III or Cd11b^+^ TC IV. Images are representative of two independent experiments with similar results. **g-h**, Immunofluorescence analysis of mLN from P17 *Rorc^Venus^* mice. Representative histo-cytometry plot for identification of RORγt^+^Foxp3^+^ pTreg cells and RORγt^+^MHCII^+^CD11c^+^CD11b^+^ TC IV (**g**) and representative immunofluorescence imaging demonstrating distribution of indicated cell types (**h**). *n* = 3 mice, 2 lymph nodes per mouse.

**Extended Data Figure 6.**
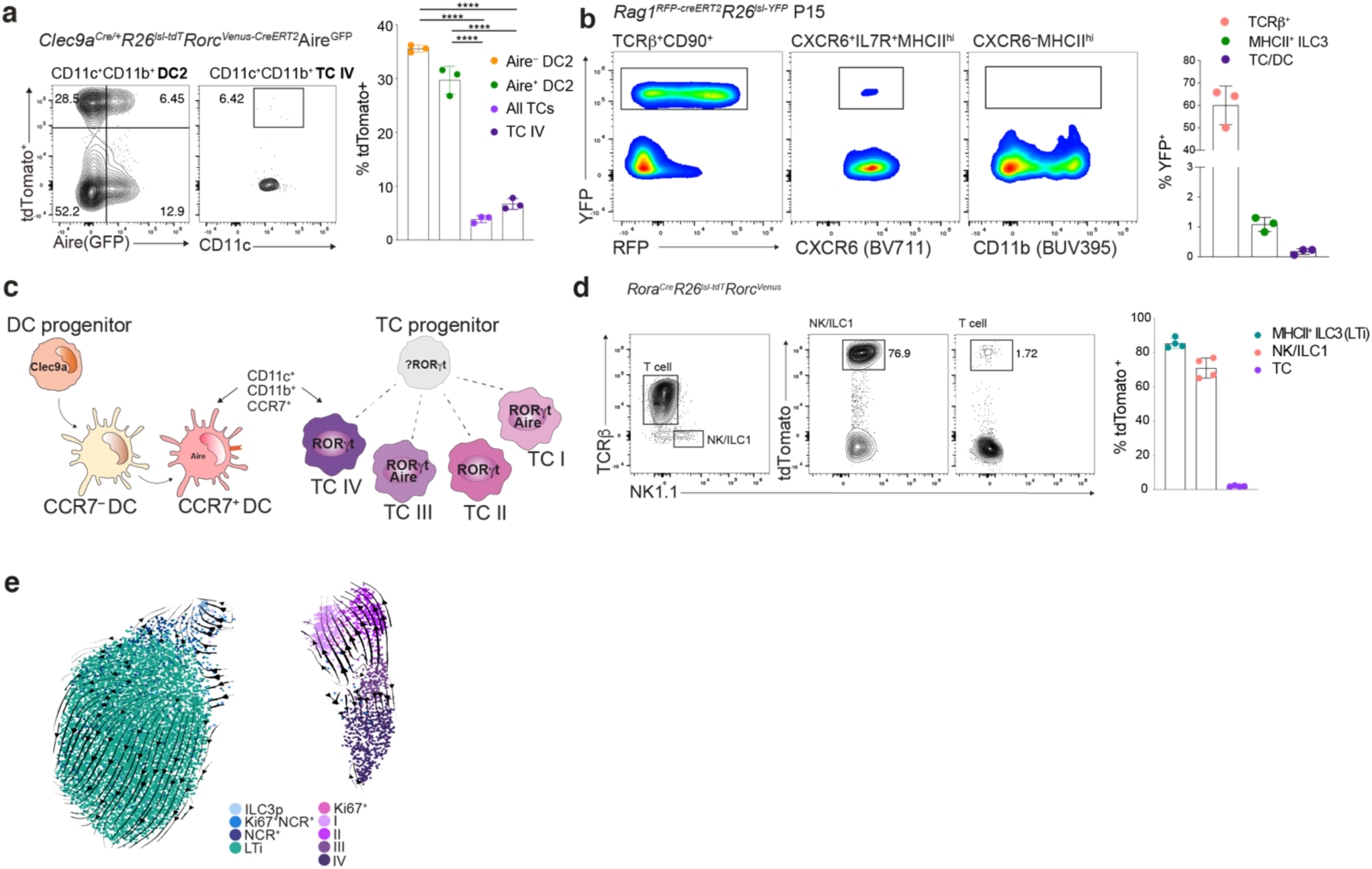
Thetis cells are ontogenically distinct from DCs and ILC3s. **a**, tdTomato labeling in cDC and TC from mLN of DC fate-mapping RORγt and Aire double reporter (*Clec9a^Cre/+^R26^lsl-tdtomato^Rorc^Venus-creERT2^*Aire^GFP^) mice at P18. **b**, Flow cytometry analysis of TCRβ^+^, MHCII^+^ ILC3, and CXCR6^−^MHCII^+^ cells encompassing TCs and DCs, from mLN of RAG1 fate-mapped (*Rag1^creERT2^R26^lsl-YFP^*) mice (*n* = 3) at P15 following 4-OHT treatment on P3, 5 and 7. **c**, Schematic of DC and TC ontogeny demonstrating distinct and overlapping transcriptional regulators and cell surface markers. **d**, Flow cytometry analysis of indicated immune cell subsets from mLN of RORα fate-mapped *Rorc^Venus^* mice and summary bar graph for tdTomato labeling. **e**, UMAP of RORγt^+^MHCII^+^ cells (Fig. 2b) with scVelo-projected velocities, shown as streamlines.

**Extended Data Figure 7.**
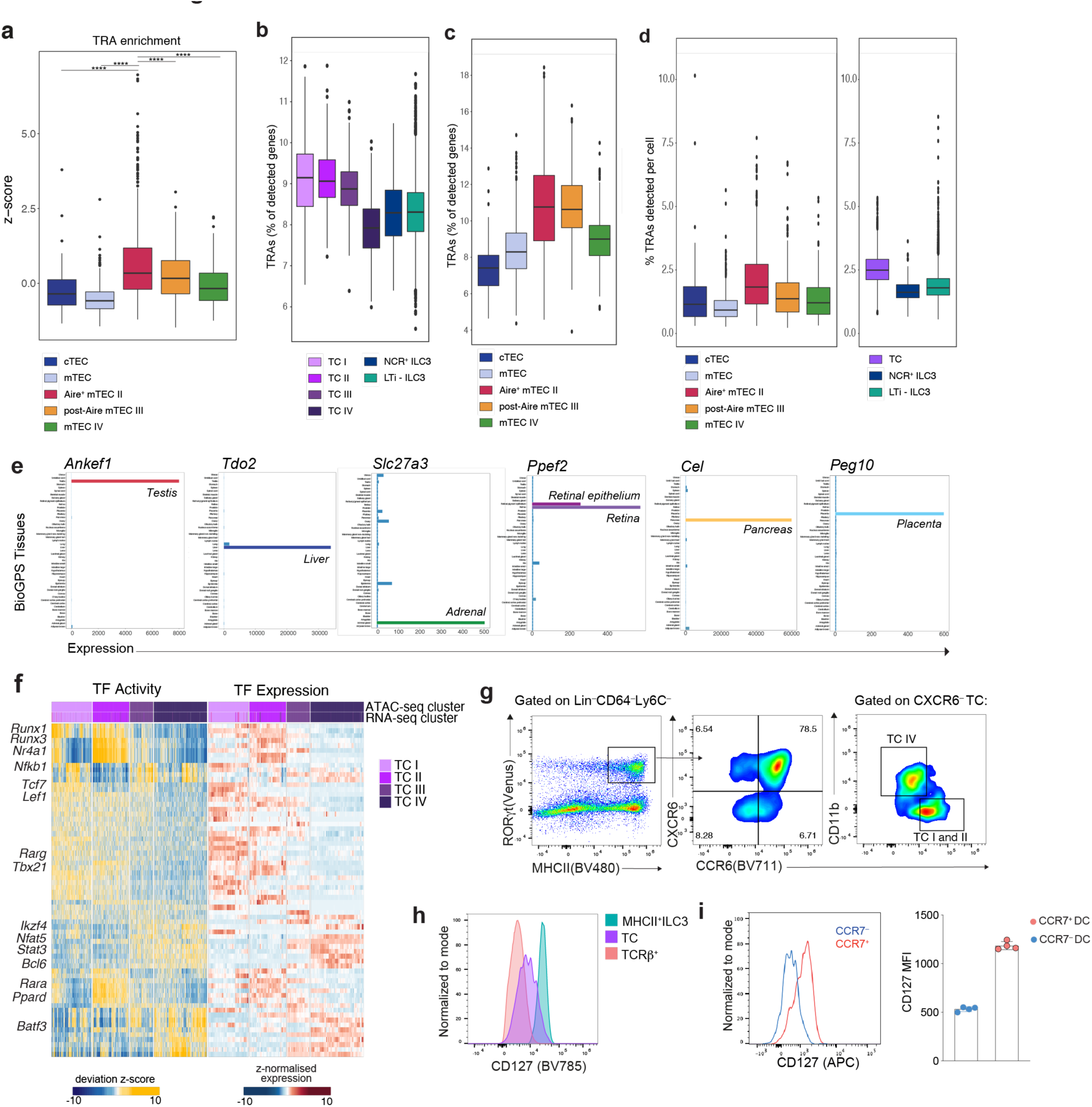
Characterization of TC subsets. **a**, Enrichment of tissue-restricted antigen (TRA) genes across published single thymic epithelial cell (TEC) transcriptomes as in Fig. 2g. **b-c**, Abundance of TRA expression by mTEC and TC subsets, normalized to the number of detected genes per cell. **d**, TRA detection rate in mTECs (left) and TCs (right). **e**, Expression pattern of example TC-enriched tissue-restricted antigens across tissues (BioGPS). **f**, Heatmap reporting TF motif activity scores for top TF-motif and gene expression pairs in scATAC/RNA-seq data (Fig. 1b). Color indicates chromVAR motif deviation score. **g**, Representative flow cytometry for CCR6 expression on ILC3 and TC subsets (*n* = 3 mice). **h**, IL7R (CD127) expression (representative of *n* = 4 mice) on ILC3s and TCs. **h**, Expression of IL7R by DC subsets. Each dot represents an individual mouse, (*n* = 4). Box plots indicate the median (center lines) and interquartile range (hinges), and whiskers represent min and max, dots represent outliers. *P*<0.0001 ****; Mann Whitney U test.

**Extended Data Figure 8.**
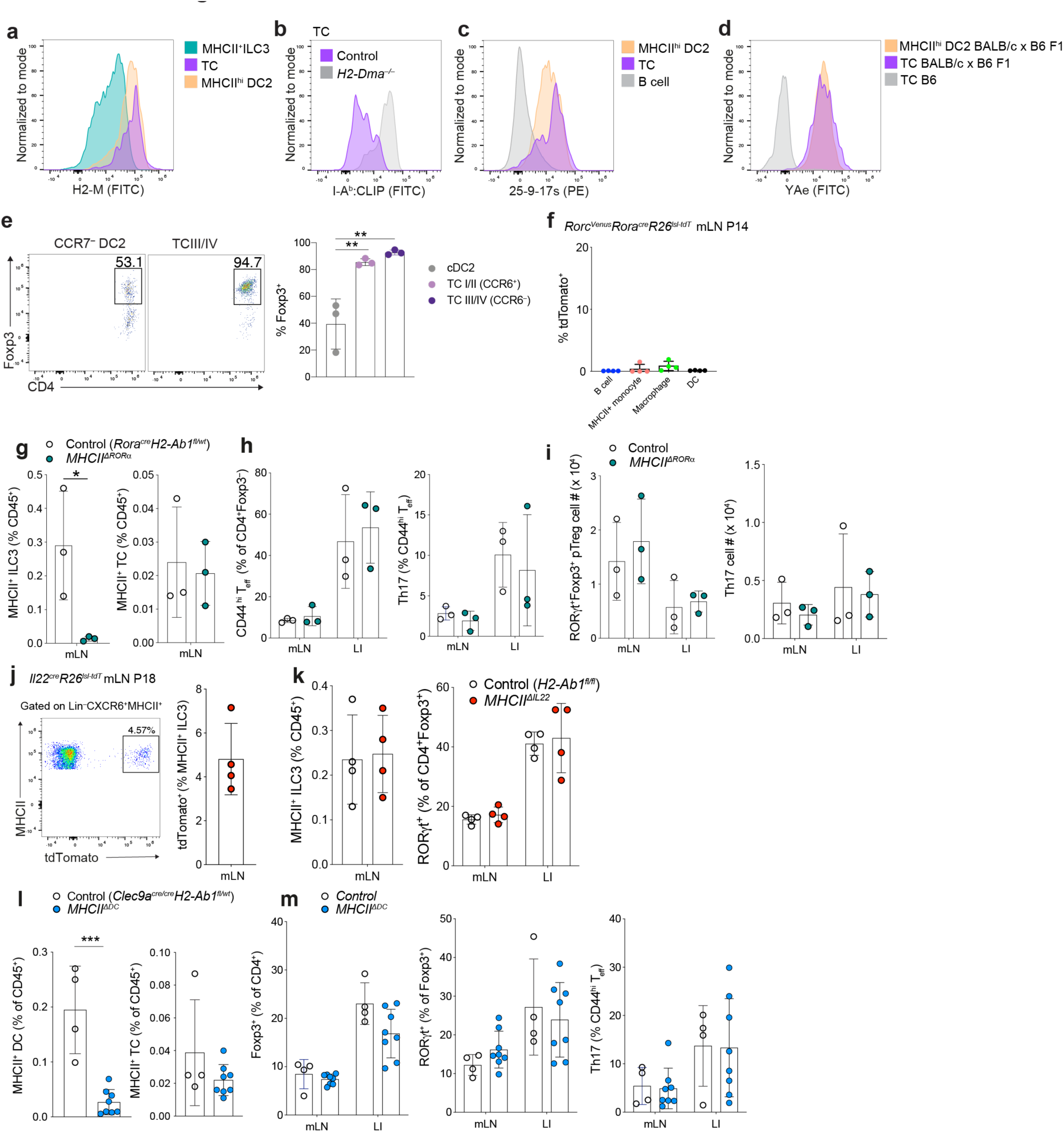
Antigen presentation by ILC3s or DCs is not required for extra-thymic intestinal pTreg differentiation. **a-d**, Flow cytometry of mLN from P14 *Rorc^Venus-creERT2^* (**a,c**) or *H2-Dma^−/−^* and littermate wild-type mice (**b**) or BALB/c x B6 F1 *Rorc^Venus-creERT2^* mice (**d**). demonstrating expression of indicated antigen processing and presenting molecules. MHCII^+^ ILC3s: (Lin^−^CXCR6^+^RORγt^+^MHCII^+^, TCs: (Lin^−^CXCR6^−^RORγt^+^MHCII^+^ and DC2s: (Lin^−^RORγt^−^CD11c^+^MHCII^hi^CD11b^+^. Representative of *n* = 4 mice. **e**, frequency of Foxp3^+^ T cells amongst CD4^+^ T cells following co-culture of naïve CD4^+^ C7 T cells with indicated TC or DC subset. **f**, tdTomato expression by MHCII^+^ cell types in mLN of *Rorc^Venus^Rora^cre^R26^lsl-tdTomato^* fate-mapped mice at P14*; n* = 4 mice. **g-i**, Immune cell composition of 3-week-old *MHCII^ΔRORα^* (*n* = 3) and *Rora*^cre^*H2-Ab1^fl/wt^* (*n* = 3) mice. (**g**) Frequency of MHCII^+^ ILC3s and TCs in mLN. (**h**), Frequency of CD4^+^Foxp3^−^CD44^hi^ T_eff_ and RORγt^+^ Th17 cells, (**i**) Total number of RORγt^+^Foxp3^+^ pTreg and RORγt^+^ Th17 cells. **j**, tdTomato labeling in MHCII^+^ILC3 (Lin^−^CXCR6^+^MHCII^+^) from mLN of IL22 fate-mapping (*Il22^Cre/+^R26^lsl-tdtomato^*) mice at P18. Representative flow cytometry and summary bar graph, n = 4 mice. **k**, Frequency of MHCII^+^ ILC3s and RORγt^+^ pTreg amongst CD4^+^Foxp3^+^ cells in indicated tissues from 3-week-old *MHCII^ΔIL22^* (*n* = 4) and control (*H2-Ab1^fl/fl^*) (*n* = 4) mice. **l-m**, Immune cell composition in mLN of 3-week-old *MHCII^ΔDC^* (*n* = 4) and control (*Clec9a*^cre^*H2-Ab1^fl/wt^*) (*n* = 8) mice from 2 independent experiments. Frequency of MHCII expressing DCs or TCs within mLN (**l**). Frequency of total Foxp3^+^ Treg cells, pTreg cells amongst CD4^+^Foxp3^+^ cells, and Th17 cells in mLN and LI (**m**). Error bars: means ± s.e.m. \*\*\**P* < 0.01, \*\*\*\**P* < 0.0001; unpaired two-sided *t*-test.

**Extended Data Figure 9.**
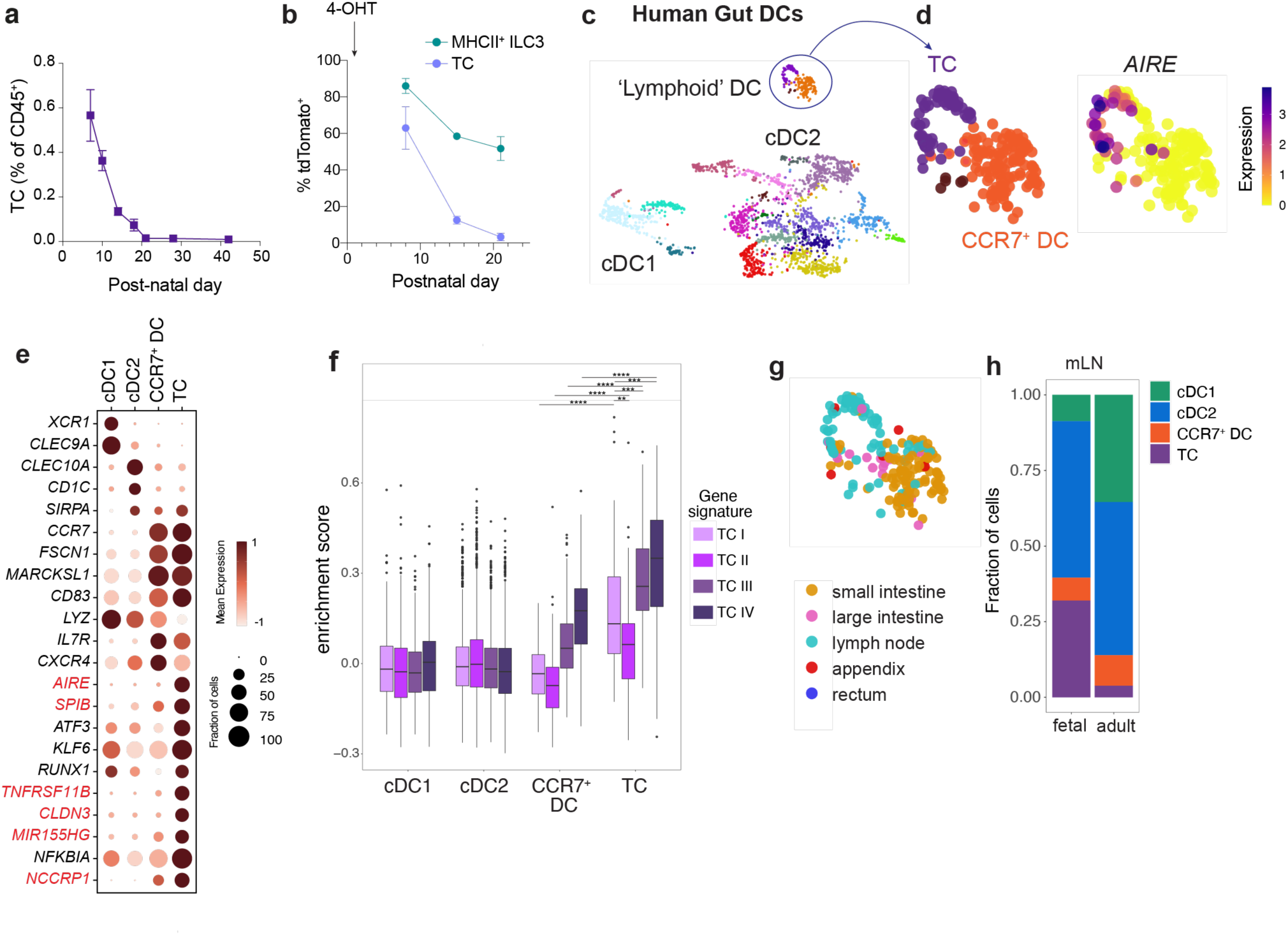
TCs are enriched in early life and conserved across mouse and human. **a**, Frequency of TCs within mLN from postnatal day 7 to 6-weeks-of age. **b**, Percentage of tdTomato^+^ TCs and MHCII^+^ ILC3s isolated from mLN of *Rorc^Venus-creERT2^R26^lsl-tdTomato^*Aire^GFP^ mice at indicated time intervals following administration of 4-OHT on P1 (*n* = 4–5 mice per timepoint). **c**, Human gut atlas single cell transcriptomes. Cells annotated as DCs were reclustered with PhenoGraph and visualized with UMAP. **d**, UMAP of ‘lymphoid’ DC clusters colored by PhenoGraph cluster or unimputed expression of AIRE. **e**, Dot plot showing select genes differentially expressed between indicated cell subsets. **f**, Enrichment of TC subset signature genes within indicated human APC subsets. **g**, UMAP colored by tissue of origin. **h**, Proportion of indicated DC/TC subsets within mLN samples in fetal *vs* adult samples. Clusters annotated as cDC2 or cDC1 were grouped for analysis. Box plots (**f**) indicate the median (center lines) and interquartile range (hinges), and whiskers represent min and max, dots represent outliers. \*\*\*\**P* < 0.0001, \*\*\**P* < 0.001, \*\**P* < 0.01; Mann Whitney U test (**f**).

**Extended Data Figure 10.**
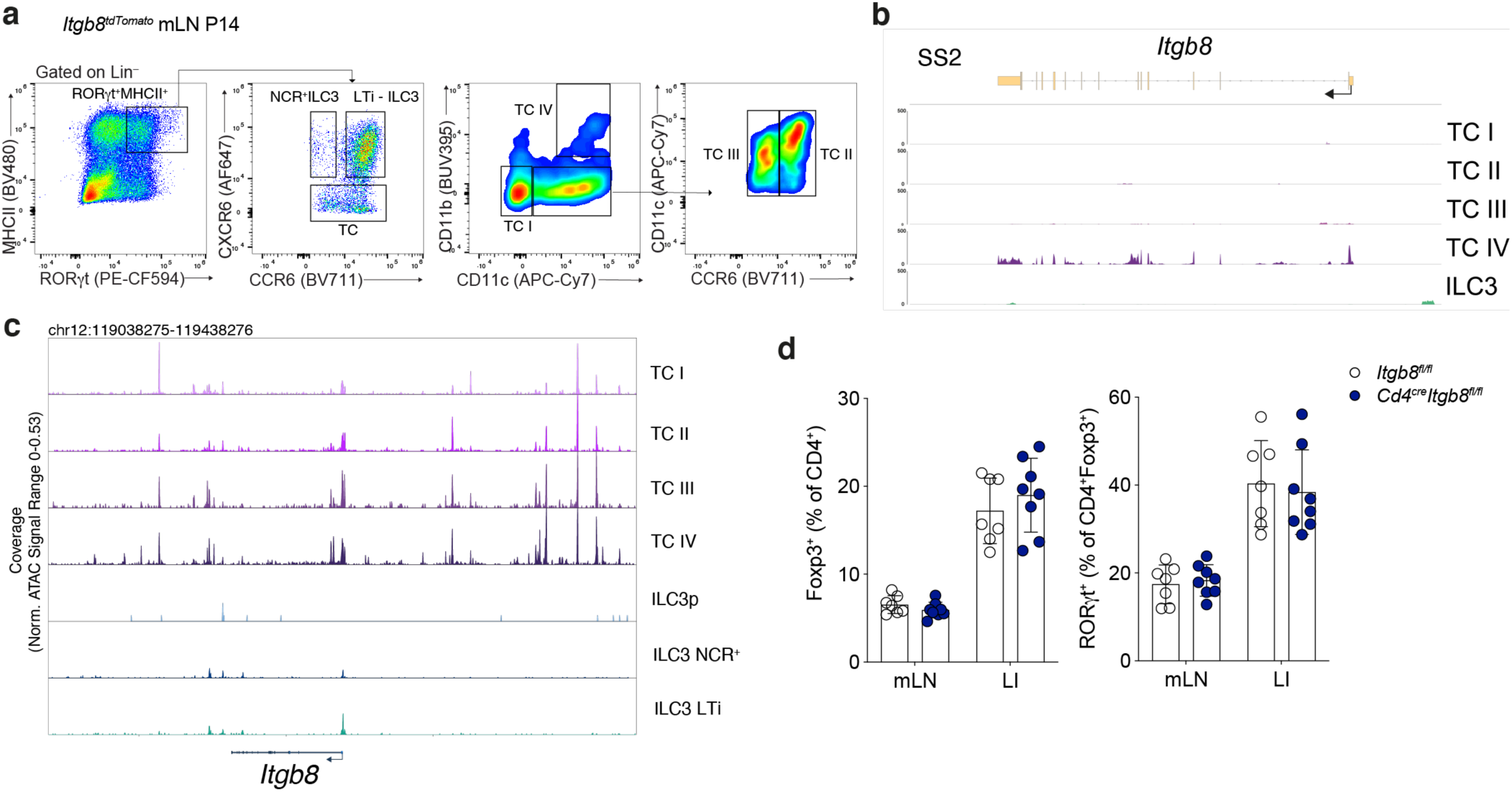
TCs promote intestinal pTreg differentiation in an Itgb8-dependent manner. **a**, Gating strategy for identification of ILC3 and TC subsets in mLN of *Itgb8^tdTomato^* mice at P14 (representative of *n* = 4 mice, three independent experiments). **b**, Itgb8 transcript levels in indicated TC and ILC3 profiled by Smart-seq 2. **c**, Chromatin accessibility at the *Itgb8* locus and Itgb8 transcript levels in TC and ILC3 subsets (cells as in Fig 1c). **d**. Frequency of total Foxp3^+^ Treg cells and percentage of RORγt^+^ pTreg cells in mLN and LI of *Itgb8^ΔCd4^* or Itgb8^fl/fl^ mice. *n* = 7 mice per group pooled from two independent experiments. Error bars: means ± s.e.m. NS, not significant; unpaired two-sided *t*-test.

